# An *in vivo* pig model for testing novel PET radioligands targeting cerebral protein aggregates

**DOI:** 10.1101/2021.12.31.473908

**Authors:** Nakul Ravi Raval, Arafat Nasser, Clara Aabye Madsen, Natalie Beschorner, Emily Eufaula Beaman, Morten Juhl, Szabolcs Lehel, Mikael Palner, Claus Svarer, Pontus Plavén-Sigray, Louise Møller Jørgensen, Gitte Moos Knudsen

**Affiliations:** Neurobiology Research Unit, Copenhagen University Hospital (Rigshospitalet), Copenhagen, Denmark; Faculty of Health and Medical Sciences, University of Copenhagen, Copenhagen, Denmark; Center for Translational Neuromedicine, University of Copenhagen, Copenhagen, Denmark; Cardiology Stem Cell Centre, Copenhagen University Hospital (Rigshospitalet), Copenhagen, Denmark; Department of Clinical Physiology, Nuclear Medicine & PET, Copenhagen University Hospital (Rigshospitalet), Copenhagen, Denmark; Department of Clinical Research, Clinical Physiology and Nuclear Medicine, University of Southern Denmark, Odense, Denmark; Department of Nuclear Medicine, Odense University Hospital, Odense, Denmark; Copenhagen Spine Research Unit, Copenhagen University Hospital (Rigshospitalet), Glostrup, Denmark

**Author notes:** **Correspondence**: Gitte Moos Knudsen, Neurobiology Research Unit, Section 8057, Rigshospitalet, Blegdamsvej 9, DK-2100 Copenhagen Ø, Denmark.

**Keywords:** Positron emission tomography, [^11^C]PIB, protein injection model, alpha-synuclein, amyloid-beta, brain imaging, autoradiography, large animal PET

## Abstract

Positron emission tomography (PET) has become an essential clinical tool for diagnosing neurodegenerative diseases with abnormal accumulation of proteins like amyloid-β or tau. Despite many attempts, it has not been possible to develop an appropriate radioligand for imaging aggregated α-synuclein in the brain for diagnosing, e.g., Parkinson’s Disease. Access to a large animal model with α-synuclein pathology would critically enable a more translationally appropriate evaluation of novel radioligands.

We here establish a pig model with cerebral injections of α-synuclein preformed fibrils or brain homogenate from postmortem human brain tissue from individuals with Alzheimer’s disease (AD) or dementia with Lewy body (DLB) into the pig’s brain, using minimally invasive surgery and validated against saline injections. In the absence of a suitable α-synuclein radioligand, we validated the model with the unselective amyloid-β tracer [^11^C]PIB, which has a high affinity for β-sheet structures in aggregates. Gadolinium-enhanced MRI confirmed that the blood-brain barrier was intact. A few hours post-injection, pigs were PET scanned with [^11^C]PIB. Quantification was done with Logan invasive graphical analysis and simplified reference tissue model 2 using the occipital cortex as a reference region. After the scan, we retrieved the brains to confirm successful injection using autoradiography and immunohistochemistry.

We found four times higher [^11^C]PIB uptake in AD-homogenate-injected regions and two times higher uptake in regions injected with α-synuclein-preformed-fibrils compared to saline. The [^11^C]PIB uptake was the same in non-injected (occipital cortex, cerebellum) and injected (DLB-homogenate, saline) regions. With its large brains and ability to undergo repeated PET scans as well as neurosurgical procedures, the pig provides a robust, cost-effective, and good translational model for assessment of novel radioligands including, but not limited to, proteinopathies.

## 1. Introduction

Several neurodegenerative diseases share the pathology of misfolded proteins (Lázaro et al., 2019), and positron emission tomography (PET) neuroimaging has become the primary imaging modality to precisely diagnose and quantify such proteinopathies in patients. As of now, many suitable PET radioligands are in use for neuroimaging of amyloid-β and tau (Mathis et al., 2017); these aggregated proteins are seen in diseases such as Alzheimer’s disease (AD), frontotemporal dementia, and progressive supranuclear palsy. By contrast, attempts to develop a suitable radioligand for neuroimaging of α-synuclein aggregates or inclusions, the hallmark of Parkinson’s disease (PD), multiple system atrophy, and dementia with Lewy bodies (DLB) have largely failed. A PET radioligand targeting α-synuclein would critically assist in an earlier and more precise diagnosis, which would be helpful for both the patient and clinician and it could facilitate development of efficacious treatments.

In preclinical studies, some radioligands have shown promise for detection of α-synuclein aggregates (Hooshyar Yousefi et al., 2019; Capotosti et al., 2020; Kaide et al., 2020; Kuebler et al., 2020), as described in an extensive review on small molecules PET imaging of α-synuclein (Korat et al., 2021). Nevertheless, because of lack of specificity or affinity to α-synuclein, no tracers have succeeded in translating to humans. α-synuclein radioligands may also require higher selectivity and affinity due to the lower aggregated protein pathology seen in α-synucleinopathies compared to the extensive pathology seen in amyloid-β- and tauopathies (Braak and Braak, 2000; Lashuel et al., 2013). Moreover, α-synuclein inclusions are mostly intracellularly located which may make them less accessible to radioligands compared to extracellular amyloid-β aggregates.

A particular challenge has been an unmet need for appropriate α-synuclein animal models. Novel PET radioligands are often initially tested in rodents due to lower costs and availability of disease models, then translated to higher species, including humans. Tracers with low rodent brain uptake are often quickly discarded although it is known that rodents have higher efflux transporter activity than larger animals (Shalgunov et al., 2020). Rodent proteinopathy models also do not entirely resemble human pathology, making it difficult to predict novel radioligands’ performance in higher species. That is, access to an appropriate large animal proteinopathy model would substantially advance the evaluation of novel radioligands for neuroimaging, e.g., α-synuclein and reduce the risk of failure due to poor translation from *in vitro* to humans. The pig has become an attractive alternative to nonhuman primates which are associated with high costs, feasibility, repeatability, and not the least, the use is associated ethical concerns (Harding, 2017).

We here propose the use of domestic pigs with intracerebral protein injections as a suitable translational model for testing new radioligands. We and others have previously made widespread use of the pig for this purpose (Parker et al., 2012; Ettrup et al., 2013; Hansen et al., 2014; Winterdahl et al., 2014; Donovan et al., 2020) because the pig has high predictive value for a successful translation to humans. In our pig model here, we make intracerebral injections of either α-synuclein preformed fibrils, postmortem AD human brain homogenate (containing amyloid-β and tau pathology), postmortem DLB human brain homogenate with pure α-synuclein pathology, and control these injections with saline. Due to the absence of an appropriate α-synuclein radioligand, we validate our model using [^11^C]PIB, a non-specific radioligand for amyloid-β (Klunk et al., 2004), which also has affinity to α-synuclein preformed fibrils but not to Lewy bodies (Ye et al., 2008). To confirm the integrity of the blood-brain barrier, we conducted gadolinium-enhanced MRI scan and to confirm brain pathology, we characterized the injected brain regions with immunohistochemistry and autoradiography.

## 2. Methods

### 2.1 Animals

Seven female domestic pigs (crossbreed of Landrace × Yorkshire × Duroc) weighing on average 27±1 kg (ranging from 25-31 kg) and approximately 10-11 weeks old were used in the present study (Table 1). Animals were sourced from a local farm and prior to any experiments. They were acclimatized for 7-9 days in an enriched environment. All animal procedures were performed following the European Commission’s Directive 2010/63/EU, approved by the Danish Council of Animal Ethics (Journal no. 2017-15-0201-01375), and complied with the ARRIVE guidelines. The overall design of the study is shown in Figure 1.

**Table 1.**
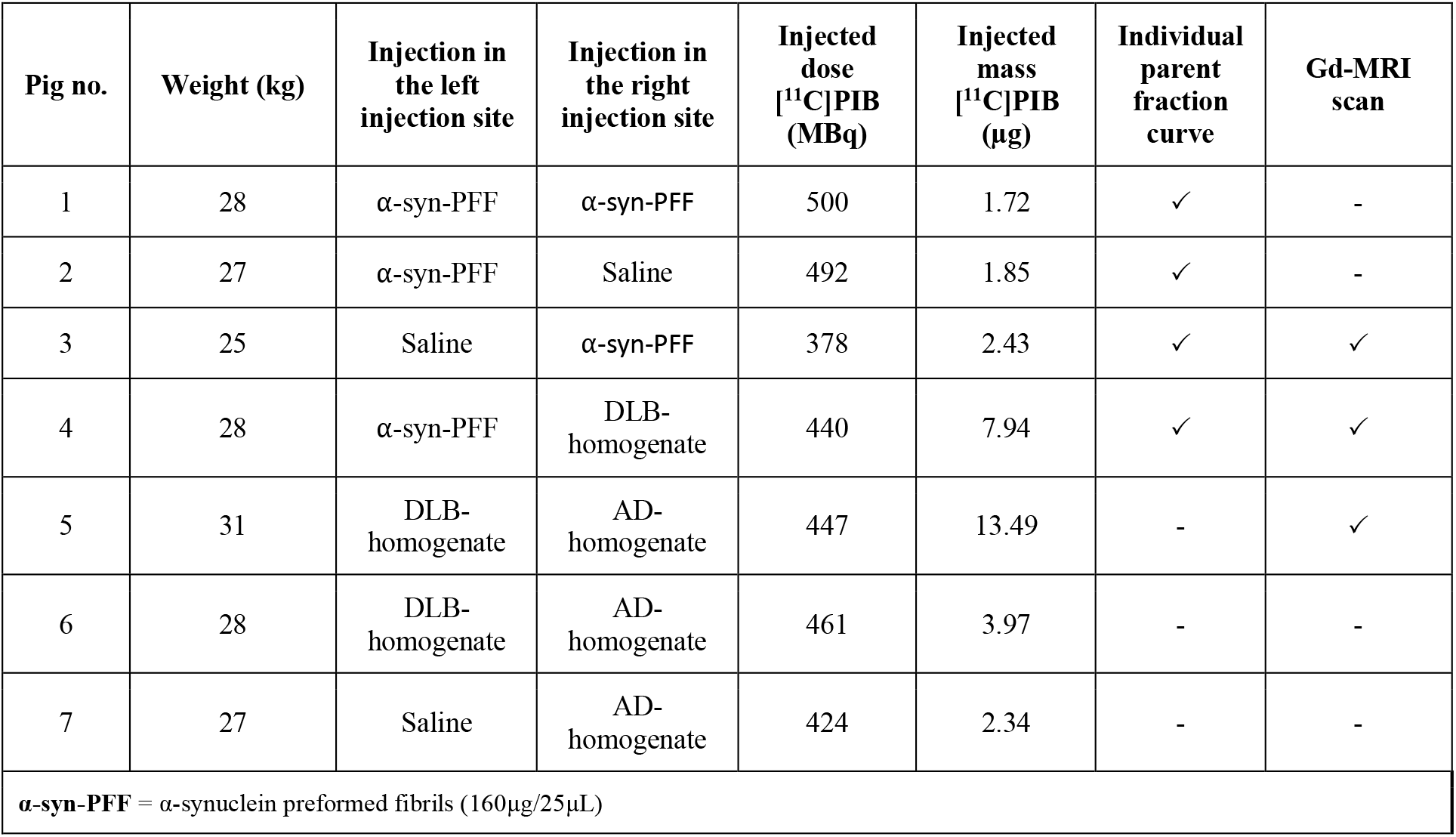
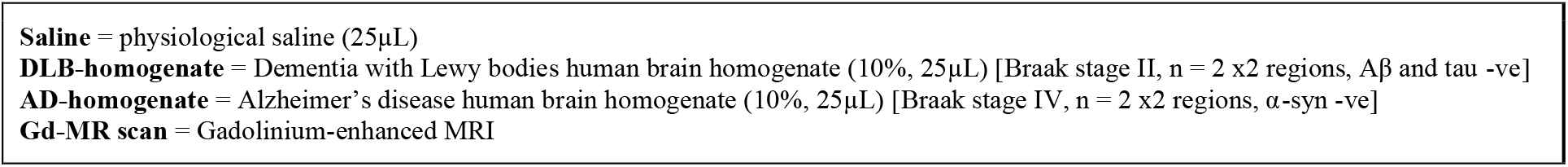
Overview of animals. Bodyweight, injection substance, injected dose/mass of [^11^C]PIB, and availability of parent fraction curve, and gadolinium contrast MR scan.

| Pig no. | Weight (kg) | Injection in the left injection site | Injection in the right injection site | Injected dose [<sup>11</sup>C]PIB (MBq) | Injected mass [<sup>11</sup>C]PIB (μg) | Individual parent fraction curve | Gd-MRI scan |
| --- | --- | --- | --- | --- | --- | --- | --- |
| 1 | 28 | α-syn-PFF | α-syn-PFF | 500 | 1.72 | ✓ | - |
| 2 | 27 | α-syn-PFF | Saline | 492 | 1.85 | ✓ | - |
| 3 | 25 | Saline | α-syn-PFF | 378 | 2.43 | ✓ | ✓ |
| 4 | 28 | α-syn-PFF | DLB-homogenate | 440 | 7.94 | ✓ | ✓ |
| 5 | 31 | DLB-homogenate | AD-homogenate | 447 | 13.49 | - | ✓ |
| 6 | 28 | DLB-homogenate | AD-homogenate | 461 | 3.97 | - | - |
| 7 | 27 | Saline | AD-homogenate | 424 | 2.34 | - | - |
| α-syn-PFF = α-synuclein preformed fibrils (160μg/25μL) | | | | | | | |

Saline = physiological saline (25µL) DLB-homogenate = Dementia with Lewy bodies human brain homogenate (10%, 25µL) [Braak stage II, n = 2 x2 regions, Aβ and tau -ve] AD-homogenate = Alzheimer's disease human brain homogenate (10%, 25µL) [Braak stage IV, n = 2 x2 regions, α-syn -ve] Gd-MR scan = Gadolinium-enhanced MRI

**Figure 1.**
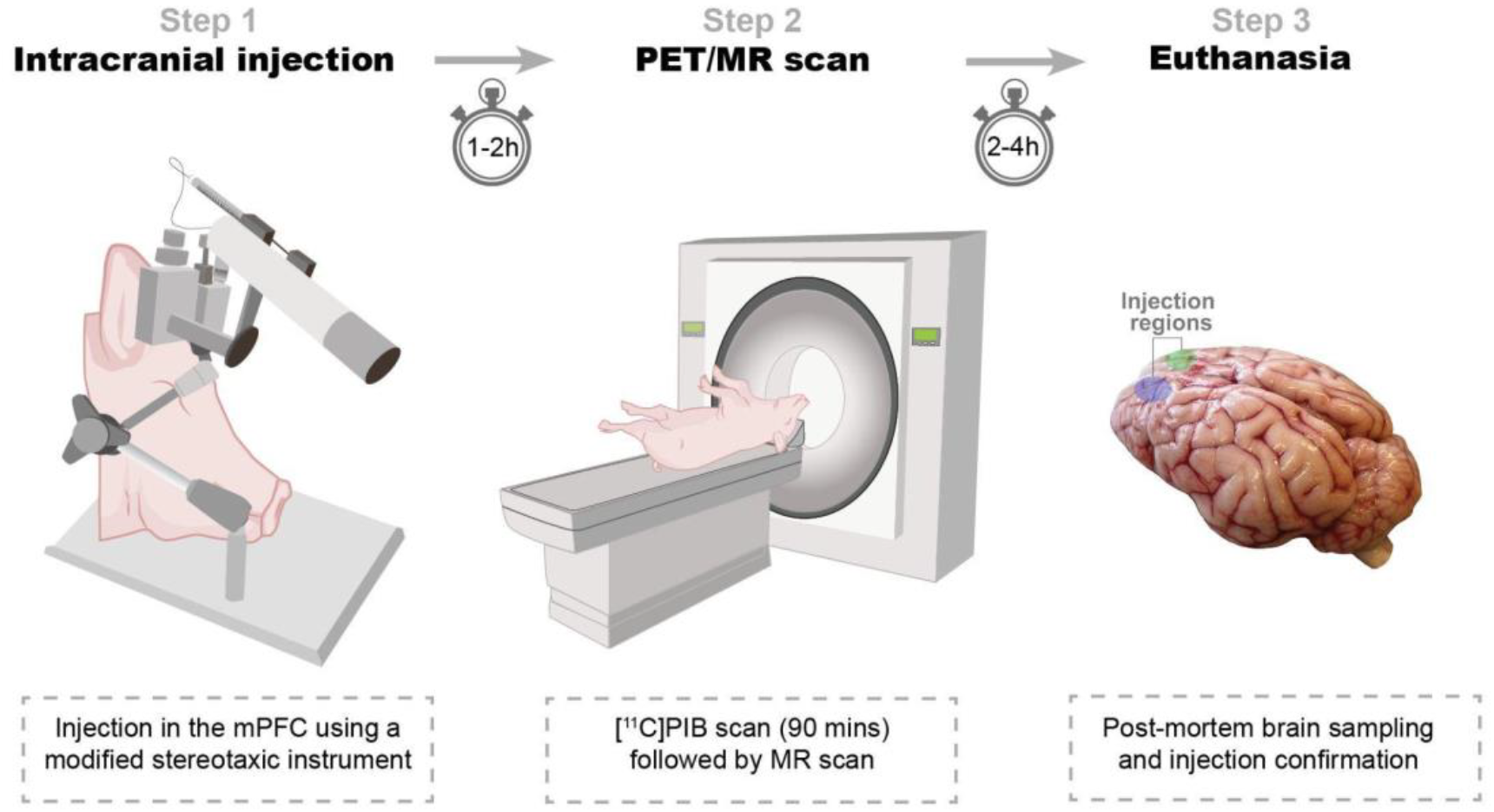
Study design. *Step 1*: Intracerebral injections. Α-synuclein preformed fibrils, Alzheimer’s disease human brain homogenate, dementia with Lewy bodies human brain homogenate, or saline is injected in either hemisphere. *Step 2*: PET/MR scan. Animals are PET scanned with [^11^C]PIB. Some animals are also MRI scanned in a 3T scanner. *Step 3*: Euthanasia. After the final scan, animals are euthanized, their brains removed, and injection sites’ pathology confirmed.

### 2.2 Preparation and surgical procedure

Pigs were injected in the medial prefrontal cortex (mPFC) with 25 μL of α-synuclein preformed fibrils (6.4 mg/mL), AD human brain homogenate (10% homogenate in saline), DLB human brain homogenate (10% homogenate in saline), or saline (Table 1). The details and characteristics of the preformed fibrils and human brains are provided in the supplementary information (Supplementary Table 1). The substrates were injected in both hemispheres, as detailed for each pig in Table 1, and in accordance with our procedure for targeting mPFC (Jørgensen et al., 2017, 2018).

A detailed description of preparation, anesthesia and transport has previously been described by us (Jørgensen et al., 2021). Briefly, anesthesia was induced by intramuscular (IM) injection of Zoletil mixture and maintained with 10-15 mg/kg/h intravenous (IV) propofol infusion. Analgesia was achieved with 5 μg/kg/h fentanyl IV infusion. Endotracheal intubation allowed for ventilation with 34% oxygen in normal air at 10-12 mL/kg. The left and right femoral arteries were catheterized with Seldinger Arterial Catheter (Arrow International, Inc., Reading, PA, USA). The left and right superficial mammary veins and ear veins were also catheterized. A urinary catheter was placed to avoid discomfort and stress throughout the surgery and scanning schedule. The animals were monitored for heart rate, blood pressure, peripheral oxygen saturation (SpO2), end-tidal CO2 (EtCO2), blood glucose, and temperature throughout the scan, except while undergoing MRI scans.

Intracerebral injections were performed using a modified stereotactic approach: An in-house instrument for modified stereotactic procedures containing a head-rest plate, a flexible arm attached with a micro-manipulator (World Precision Instruments, Sarasota, FL, USA), and a micro-syringe infusion pump system (World Precision Instruments, Sarasota, FL, USA) (Supplementary Figure 1). The flexible arm allowed the micro-manipulator to be positioned and fixed relative to the target entry point with a trajectory perpendicular to the skull, as illustrated in Supplementary Figure 1. For the first two experiments (Pig 1 and 2), we used a prototype of the device with slightly less arm flexibility and a different micro-manipulator brand and syringe-type, although the capacity, needle size, length, and tip shape were the same. However, the prototype did provide injections comparable to the remaining ones, as validated with immunohistochemistry.

After installation of local anesthesia, midline incision, and skull exposure, two burr holes were placed bilaterally, 25 mm anterior and 8 mm lateral to bregma, followed by hemostasis and dura puncture. We have previously validated this target point: 8, 25, 14 mm in the X, Y, Z coordinate relative to bregma, to center on grey matter in the mPFC (Jørgensen et al., 2017, 2018). The syringe and the needle were then positioned and fixed in a trajectory perpendicular to the skull and with the needle tip adjusted to the skull entry point. The syringes (250 μL SGE Gas-tight Teflon Luer Lock Syringes (World Precision Instruments, Sarasota, FL, USA) [different syringes for the different injectates]) were attached with SilFlex tubing (World Precision Instruments, Sarasota, FL, USA), NanoFil Injection Holder (World Precision Instruments, Sarasota, FL, USA) and 28 G Hamilton Kel-F hub blunt tip needle (Hamilton Central Europe, Giarmata, Romania). The SilFlex tubing and NanoFil Injection Holder were removed during homogenate injection because of the viscous content.

Using the micromanipulator, the needle was slowly advanced to the mPFC target point (perpendicular to the skull). The injection was performed over two steps with 10 μL, and 15 μL injected 1 mm apart (centered at the mPFC target point). The infusion was delivered at 450 nL/min using the micro-syringe infusion pump followed by a 7-minute pause before a slow withdrawal of the needle to avoid backflow. After the procedure, both burr holes were packed with an absorbable hemostatic gelatin sponge (Curaspon^®^, CuraMedical BV, Assendelft, Netherlands), and the incision was sutured shut. The animals were then transported to the scanner facilities and connected to the respirator.

### 2.3 PET scanning protocol and radiochemistry

All pigs were PET-scanned with a Siemens high-resolution research tomograph (HRRT) scanner (CPS Innovations/Siemens, Malvern, PA, USA). [^11^C]PIB was prepared at the Copenhagen University Hospital, Rigshospitalet, as per routine clinical preparation. The complete method of preparation is explained in the supplementary information (Supplementary Figure 2). Data acquisition lasted 90 min after bolus injection (over ~20 s) of [^11^C]PIB through one of the superficial mammary veins (IV). The injected dose was 448±41 MBq, while injected mass was 4.82±4.3 μg (mean±SD).

### 2.4 Blood sampling and HPLC analyses

Manual arterial blood samples were drawn at 2.5, 5, 10, 20, 30, 40, 50, 70, and 90 min after injection, while an ABSS autosampler (Allogg Technology, Strängnäs, Sweden) continuously measured arterial whole blood radioactivity during the first 20 min. The manual blood samples were measured for total radioactivity in whole blood and plasma using an automated gamma counter (Cobra 5003; Packard Instruments, Downers Grove, CT, USA) cross-calibrated against the HRRT. Radiolabeled parent and metabolite fractions were determined in plasma using an automatic column-switching radio-high performance liquid chromatography (HPLC) as previously described (Gillings, 2009), equipped with an extraction column Shim-pack MAYI-ODS (50 μm, 30 x 4.6 mm; Shimadzu Corporation, Kyoto, Japan) eluting with 50 mM HNa_2_PO_4_ pH 7.0 and 2% isopropanol (v/v) at a flow rate of 3 mL/min, and an Onyx Monolithic C18 analytical column (50 x 4.6 mm, Phenomenex, Torrance, CA, USA) eluting with 26% acetonitrile and 74% 100 mM HNa_2_PO_4_ pH 7.0 (v/v) at a flow rate of 3 mL/min. Before analysis by radio-HPLC, the plasma samples were filtered through a syringe filter (Whatman GD/X 13 mm, PVDF membrane, 0.45 m pore size; Frisenette ApS, Knebel, Denmark). Plasma was diluted 1:1 with the extraction buffer, and up to 4 mL of plasma sample was used. The eluent from the HPLC system was passed through the radiochemical detector (Posi-RAM Model 4; LabLogic, Sheffield, UK) for online detection of radioactive parent and metabolites. Eluents from the HPLC were collected with a fraction collector (Foxy Jr FC144; Teledyne, Thousand Oaks, CA, USA), and fractions were counted offline in a gamma well counter (2480 Wizard2 Automatic Gamma Counter, PerkinElmer, Turku, Finland). The parent fraction was determined as the percentage of the radioactivity of the parent to the total radioactivity collected. Examples of radio-HPLC chromatograms from a pig are shown in Supplementary Figure 3.

### 2.5 Gadolinium-contrast MRI scanning protocol

The integrity of the BBB post-intracerebral injection was assessed by determining the %-difference ΔT1-map of the pre-gadolinium and the post-gadolinium scans. The MRI data were acquired on a 3T Prisma scanner (Siemens, Erlangen, Germany) using a human 64-channel head coil (active coil elements were HC3-7 and NC1). Three pigs were scanned in the MRI scanner as previously described by us (Jørgensen et al., 2021). The pigs underwent two T1-map scans: pre-and post-gadolinium contrast injection. The protocol for T1-weighted 3D magnetization-prepared rapid gradient-echo (MP-RAGE) MRI was: frequency direction = anterior to posterior; dimension= 144 x 256 x 256; slice thickness = 0.9 mm; repetition time = 2000 ms; echo time = 2.58 ms; inversion time = 972 ms; flip angle = 8°; base resolution = 256 x 256, and acquisition time = 192 s. After the pre-gadolinium T1-map scan, pigs received gadolinium IV (0.1 mmol/kg, Gadovist® [gadobutrol], Bayer A/S, Copenhagen, Denmark) through a superficial mammary vein and were rescanned 5 mins later with another T1-map scan. Data were processed using a custom code in MATLAB 9.5.0 (R2018b) (The MathWorks Inc., Natick, MA, USA). DICOM files were converted to NIfTI-1 using dcm2niix (Li et al., 2016). The post-gadolinium T1-map was co-registered and resliced to the pre-gadolinium T1-map using SPM12. A %-difference map (ΔT1-map) was created from the resliced post-gadolinium and pre-gadolinium T1-maps (Equation 1). Three regions in the ΔT1-map were measured: left injection site, right injection site, and occipital cortex

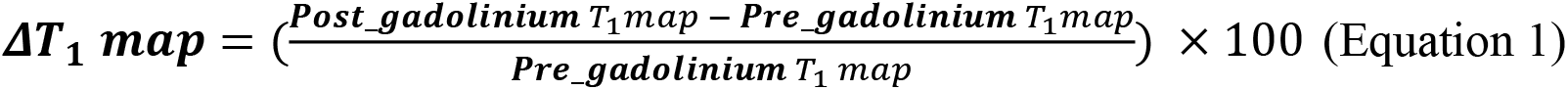

### 2.6 [^3^H]PIB autoradiography

At the end of the scanning, the animals were euthanized by IV injection of 20 mL pentobarbital/lidocaine. After euthanasia, the brains were removed, snap-frozen with powdered dry-ice, and stored at −80°C until further use. 20 μm coronal cryosections were sectioned on a cryostat (Thermo Scientific/Epredia™ CryoStar™ NX70 Cryostat, Shandon Diagnostics Limited, Runcorn, UK) and mounted on Superfrost Plus™ adhesion microscope slides (Thermo Fisher Scientific, Waltham, MS, USA). Sections were stored at −80°C for the remaining period of the study.

We performed [^3^H]PIB (Novandi Chemistry AB, Södertälje, Sweden, Molar activity: 78 Ci/mmol) autoradiography to calculate the total available binding sites (*B*_max_) and equilibrium dissociation constant (*K*_D_) in the injected pig brain, and compared this to the *B*_max_ and *K*_D_ of human brain regions that were used to create the homogenates. We performed saturation assays using increasing concentrations of [^3^H]PIB for total binding and [^3^H]PIB + thioflavin S (100 μM) for non-specific binding on AD-homogenate-injected pig brain slices (n=1, 0 to 5 nM of [^3^H]PIB), α-synuclein-preformed-fibril-injected pig brain slices (n=1, 0 to 5 nM of [^3^H]PIB), and AD post-mortem human brain slices (n=2 x 2 regions, 0 to 40 nM of [^3^H]PIB). Since there was no specific binding in DLB-homogenate-injected pig slices or DLB human brain slices, we could not perform saturation assays on these sections. Human brain slices were prepared in the same fashion as pig brain slices, including section thickness and storage. Detailed procedure for autoradiography is available in the supplementary information.

The data were analyzed using GraphPad Prism (v. 9.2.0; GraphPad Software, San Diego, CA, USA). Non-linear regression analysis (Function: One site - total and non-specific binding) was used to calculate *B*_max_ and *K*_D_ values for all three assays. The fitting method used was the least squared regression with no weighting. In vitro binding potential (BP) was calculated with Equation 2.

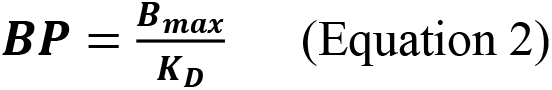

### 2.7 PET data reconstruction and preprocessing

PET list-mode emission files were reconstructed using the OP-3D-OSEM algorithm, including modeling the point-spread function, with 16 subsets, ten iterations, and standard corrections (Sureau et al., 2008). During reconstruction, attenuation correction was performed using the MAP-TR μ-map (Keller et al., 2013). Emission data were binned into time frames of increasing lengths: 6 × 10 s, 6 × 20 s, 4 × 30 s, 9 × 60 s, 2 × 180 s, 8 × 300 s, 3 × 600 s. Each time frame consisted of 207 planes of 256 × 256 voxels of 1.22 × 1.22 × 1.22 mm in size.

Brain parcellation was done with our previously published automatic PET-MR pig brain atlas method (Villadsen et al., 2017). The neocortex, occipital cortex, and cerebellum non-vermis (henceforth denoted as the cerebellum) were extracted from the Saikali atlas (Saikali et al., 2010) for the present study. Two additional regions for the injection site were hand-drawn on the atlas from an approximate injection site that was initially characterized around the site of needle penetration as visualized by the MRI scans and postmortem extracted brain. This was also further confirmed and optimized by positive immunohistochemistry slices from the region (Supplementary Figures 4 and 5). Regions approximately 0.32-0.35 cm^3^ (~ 250 voxels) in size were placed symmetrically in the left and right hemispheres. This region is slightly larger than the injection site itself, but this gives leeway for any potential mechanical error during the stereotactic operation. Wherever possible (not possible in, e.g., saline-injected regions), the automatic region was visually inspected with late-scan frames averaged.

Regional radioactivity concentration (kBq/mL) was normalized to injected dose (MBq) and corrected for the animal weight (kg) to provide standardized uptake values (SUV, g/mL) used to make average plots as in Figure 2. PMOD 3.7 (PMOD Technologies, Zürich, Switzerland) was used to visualize and create the representative PET and MR images.

**Figure 2.**
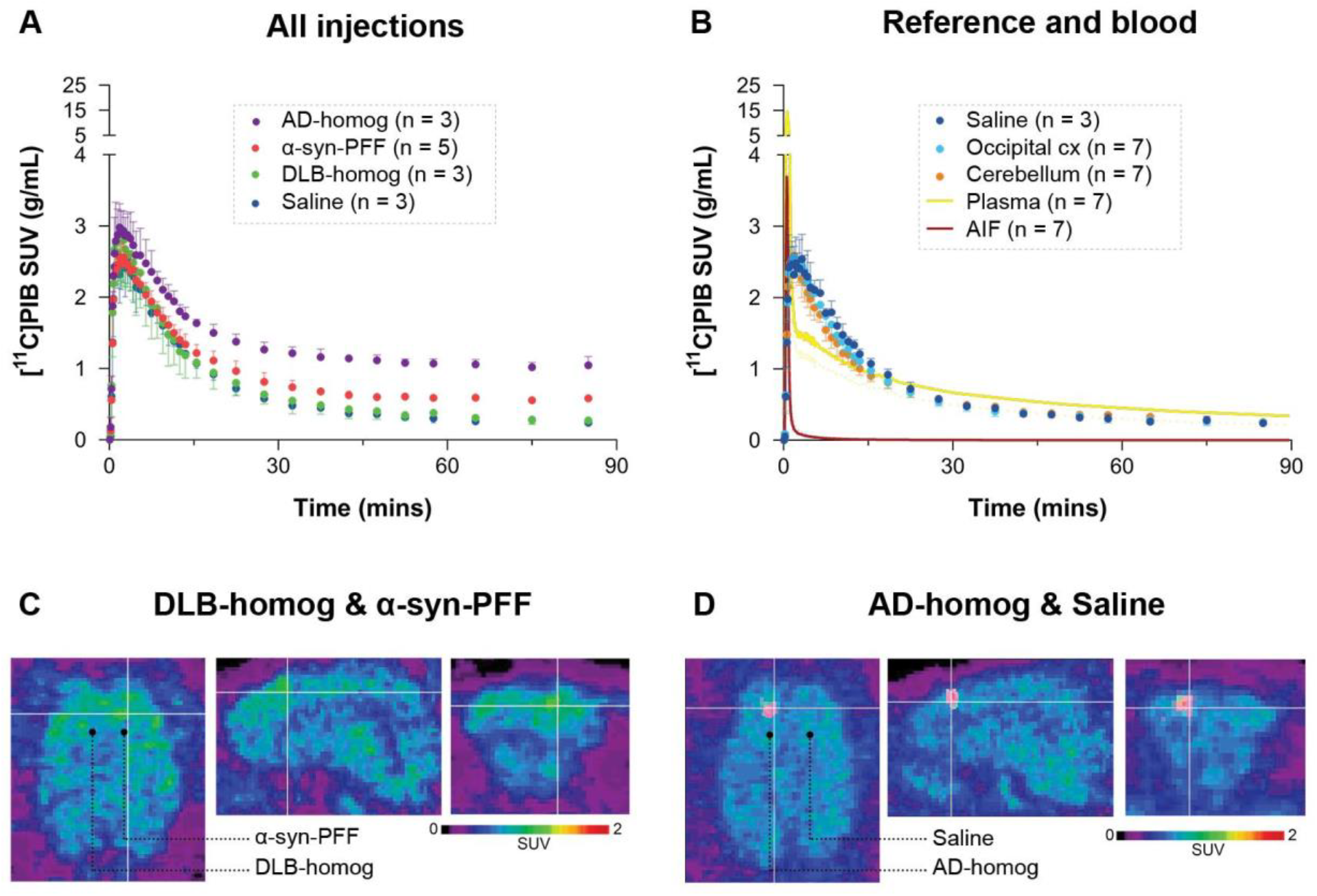
Regional time-activity curves of [^11^C]PIB in A) the different injection regions and B) the reference regions and saline-injected region with the uncorrected plasma curve and arterial input function. Representative summed PET across the entire duration of the scan (0-90 mins) images showing injection regions including C) SUV scaled brain images including the brain areas injected with α-synuclein preformed fibrils or DLB homogenate and D) AD homogenate or saline.

### 2.8 Kinetic modeling

Kinetic modeling was performed using *kinfitr* (v. 0.6.0) (Matheson, 2019; Tjerkaski et al., 2020) in R (v. 4.0.2; “Taking Off Again,” R core team, Vienna, Austria). The Logan Graphical Analysis was applied to estimate the total volume of distribution (V_T_) values (Logan et al., 1990), using a metabolite corrected input function derived from radioactivity measurements of arterial blood samples. Reference tissue modeling was performed with the simplified reference tissue model 2 (SRTM2), with an average k_2_’, to calculate non-displaceable binding potential (BPND) using the occipital cortex as the reference region (Yaqub et al., 2008). For more details on the kinetic modeling, see supplementary information.

### 2.9 Statistical analyses

Graph-Pad Prism (v. 9.2.0; GraphPad Software, San Diego, CA, USA) was used for data visualization and statistical analysis. All data are presented as mean values ± standard deviation. The difference in PET outcomes (Logan V_T_ and SRTM2 BPND) between the injected regions and reference tissues was calculated using the non-parametric Kruskal–Wallis one-way analysis of variance (ANOVA). For assessment of change in gadolinium-contrast MR, we used the Friedman non-parametric ANOVA test with paired testing. Post-hoc ANOVA tests were corrected for multiple comparisons by Dunn’s multiple comparison test (Dunn, 1964).

## 3 Results

### 3.1 [^11^C]PIB time-activity curves

After [^11^C]PIB injection, we observed high brain uptake and rapid tracer wash-out. The blood and brain kinetics of [^11^C]PIB were very fast, with less than 10% of the parent radioligand remaining in plasma after 2.5 mins (Figure 2 and Supplementary Figure 6). We found higher radioactivity retention in the AD-homogenate- and α-synuclein-preformed-fibrils injected region (Figure 2A). Compared to the cerebellum and the occipital cortex, almost no retention was seen in DLB-homogenate and saline-injected regions (Figure 2A and B).

### 3.2 Kinetic modeling of [^11^C]PIB

[11C]PIB binding parameters from Logan graphical analysis and SRTM2 are summarized in Table 2. We found 4-fold higher V_T_ values in the AD-homogenate injected region compared to the occipital cortex (p = 0.006) and 2-fold higher V_T_ values in the α-synuclein-preformed fibrils region (p = 0.034) (Figure 3A). We found no difference between the saline- and DLB-injected regions.

**Table 2.** Summary of kinetic modeling outcomes of [^11^C]PIB. All values denote the mean ± standard deviation. PFF=preformed fibrils. NA= not applicable.

| Regions | Kinetic Modelling Outcome |  |
| --- | --- | --- |
| | $V_T$<br>(ml/cm<sup>3</sup>) | $BP_{ND}$ |
| α-syn-PFF | 47.7 ± 6.3 | 0.65 ± 0.18 |
| AD-homogenate | 118.1 ± 12.9 | 2.34 ± 0.31 |
| DLB-homogenate | 31.1 ± 6.1 | 0.11 ± 0.16 |
| Saline | 21.2 ± 6.1 | 0.05 ± 0.03 |
| Occipital cortex | 23.8 ± 5.5 | NA (reference) |
| Cerebellum | 25.8 ± 6.8 | 0.01 ± 0.03 |

**Figure 3.**
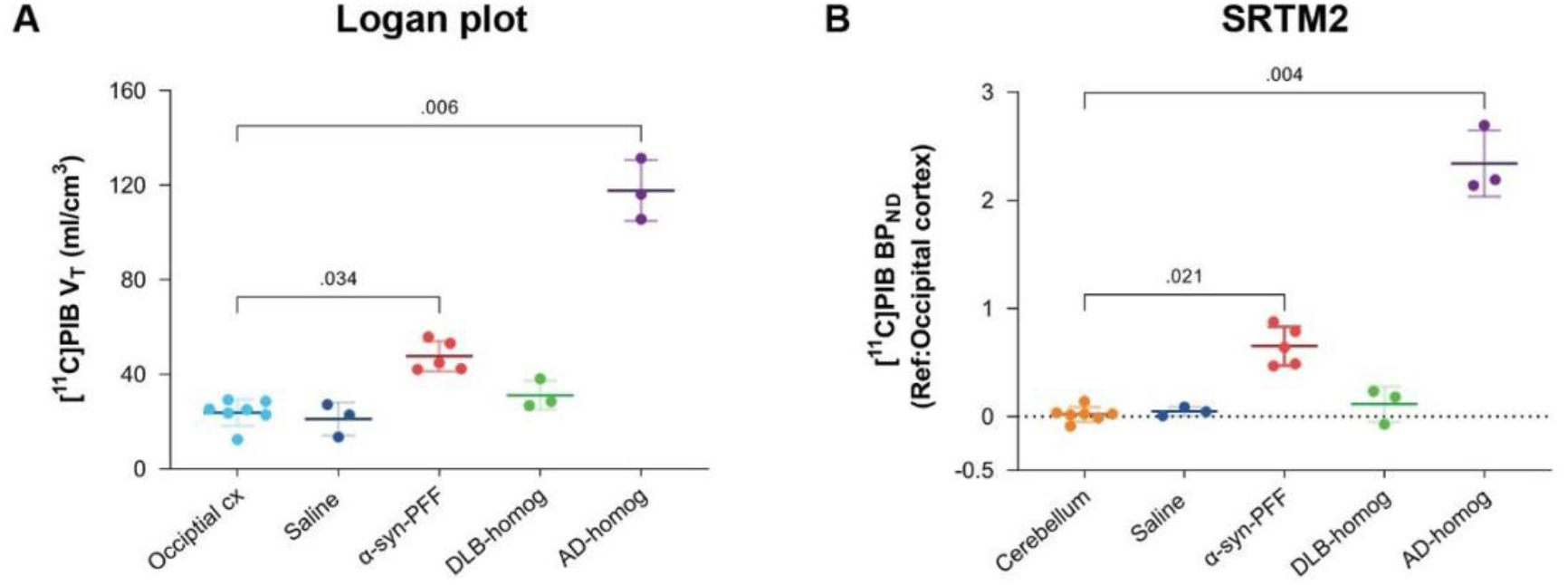
Kinetic modeling of [^11^C]PIB. A) Kinetic modeling with arterial input (Logan). Direct comparison of V_T_ values in the different injection regions to the occipital cortex. B) Kinetic modeling with occipital cortex as a reference region (SRTM2). BP_ND_ values are compared to the cerebellum.

Compared to the cerebellum, the average BP_ND_ of 2.34 was higher (p = 0.004) in the AD-homogenate injected region, and the average BP_ND_ of 0.65 was also higher (p=0.016) in the α-synuclein-preformed-fibrils injected region. There was no difference in BP_ND_ in the saline- or DLB-homogenate injected regions compared to the cerebellum (Figure 3B).

### 3.3 Characterization of the injection site

Using our minimally invasive method, we successfully injected all animals in the same symmetrical brain region. Prefrontal cortical immunostaining (α-synuclein and amyloid-β) and thioflavin S staining at the injection site confirmed the presence of α-synuclein preformed fibrils, AD homogenates, and DLB homogenates, respectively (Supplementary information). To evaluate the appropriateness of our pig model, we compared *B*_max_ and *K*_D_ in both the pig and human brains. This was done for the α-synuclein-preformed fibrils and AD-homogenate injected pig brain regions as well as for the AD postmortem human brain slices using [^3^H]PIB autoradiography saturation assays (Figure 4 and Table 3). We determined *B*_max_ to be 477.2 fmol/mg TE and *K*_D_ of 12.07 nM in the α-synuclein-preformed fibrils region in pig brain slices (n=1). We found a higher *B*_max_ on the AD postmortem human brain slices (366.7 fmol/mg TE, n=3) compared to AD-homogenate-injected pig brain slices (233.4 fmol/mg TE, n=1). However, the *K*_D_ is similar at 2.46 nM in AD-homogenate-injected pig brain slices versus 2.54 nM in AD postmortem human brain slices. We also found 2.4 times higher binding potential in AD-homogenate-injected pig brain slices compared to α-synuclein-preformed fibrils-injected pig brain slices.

**Figure 4.**
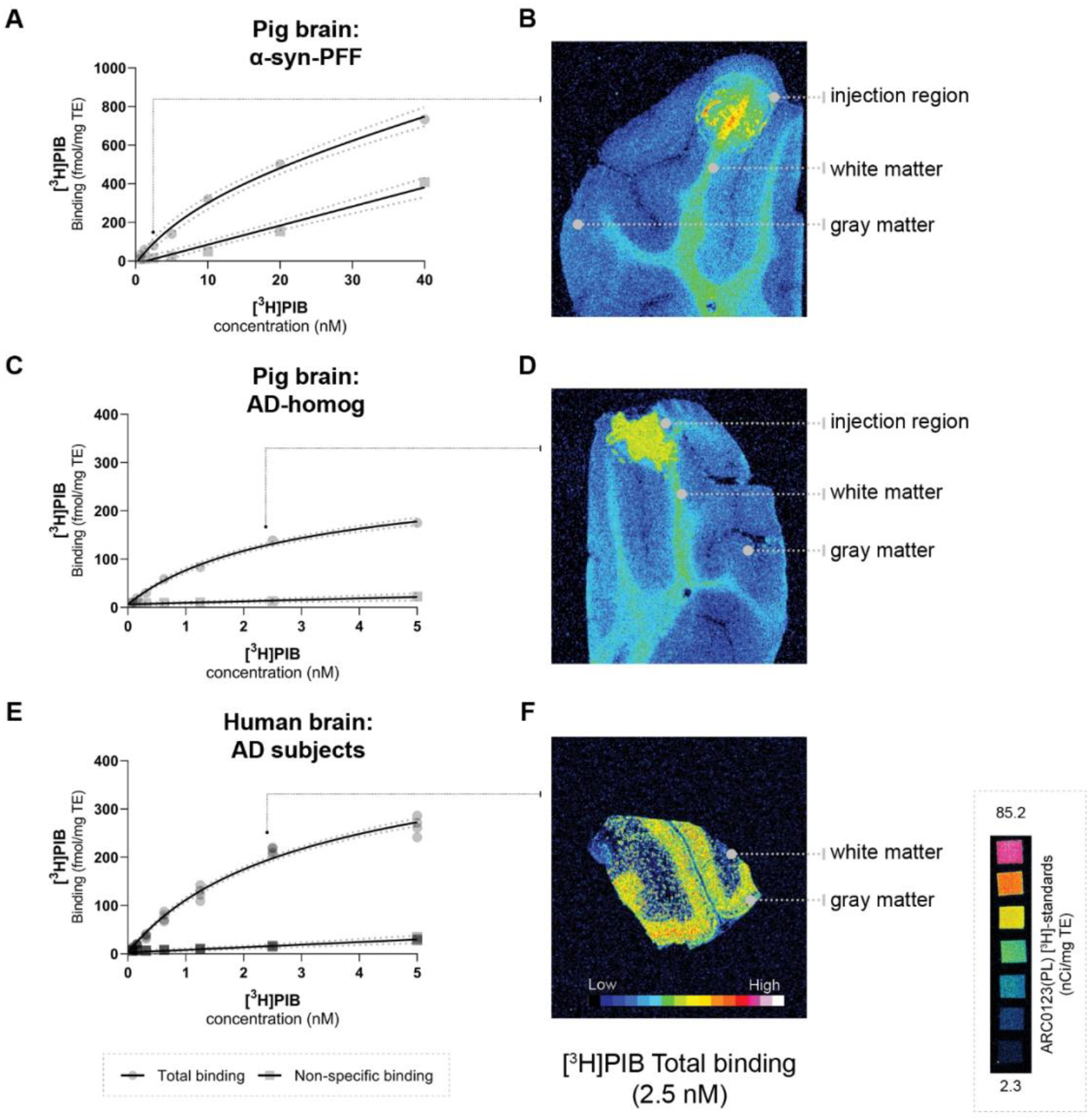
Saturation assays (A, C, E) and: corresponding representative autoradiograms (B, D, F [total binding at 2.5 nM]) of [^3^H]PIB in the pig brain: A: ⍺-syn-PFF injected, D: AD-homogenate injected, and F) Human AD brain. Scale (ARC0123(PL)) inserted.

**Table 3.**
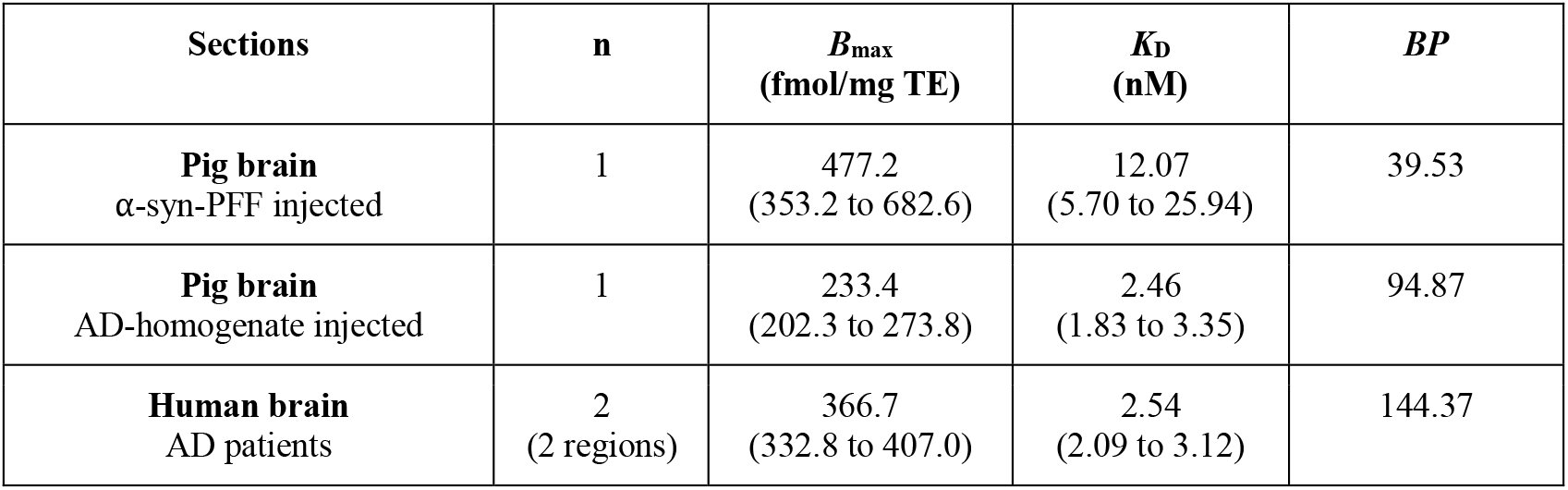
Summary of *B*_max_ and *K*_D_. Values (95% confidence interval) from [^3^H]PIB saturation assays performed on injected pigs and humans frozen brain sections. n= number of unique individuals. BP= binding potential.

| Sections | n | $B_{\max}$<br>(fmol/mg TE) | $K_D$<br>(nM) | $BP$ |
| --- | --- | --- | --- | --- |
| **Pig brain**<br>α-syn-PFF injected | 1 | 477.2<br>(353.2 to 682.6) | 12.07<br>(5.70 to 25.94) | 39.53 |
| **Pig brain**<br>AD-homogenate injected | 1 | 233.4<br>(202.3 to 273.8) | 2.46<br>(1.83 to 3.35) | 94.87 |
| **Human brain**<br>AD patients | 2<br>(2 regions) | 366.7<br>(332.8 to 407.0) | 2.54<br>(2.09 to 3.12) | 144.37 |

### 3.4 Blood-brain barrier integrity

We found no statistically significant difference in the T_1_-maps before and after gadolinium injection. Still, in cases, with local hemorrhage near the site of needle penetration (Figure 5A, red ROI), some regions had higher gadolinium uptake than the occipital cortex. Also, the amplitude of the [^11^C]PIB time-activity curves (Figure 2B and Figure 3) did not suggest that the injected regions had higher uptake compared to non-injured brain tissue. Finally, the uptake in saline-injected regions did not differ from that of the occipital cortex, supporting that the injection itself does not hamper the integrity of the BBB.

**Figure 5.**
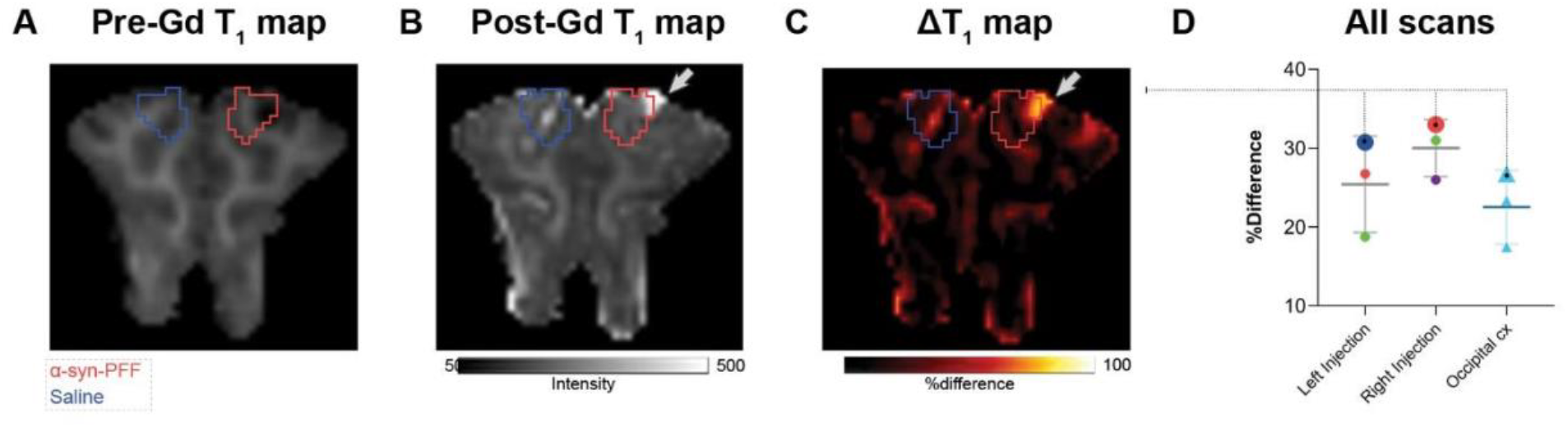
Representative pre- (A) and post- (B) gadolinium-enhanced MRI of the injected region. ΔT1 maps are shown as %difference, i.e. % (post-Gd - pre-Gd) / pre-Gd. The right and left injection regions (Right Injection vs. Left Injection) were compared to the occipital cortex. Data points were color-coded for the different injections with larger symbols from the animals shown in A-C: red circles = α-synuclein preformed fibrils injected region, dark blue circle = saline injected region, green circles = DLB homogenate injected region, purple circle = AD homogenate injected region, and light blue triangles = occipital cortex.

## 4 Discussion

We here describe a large animal model for testing radioligands against specific targets, such as abnormally configured protein structures, and the study is built on amyloid-β and its radiotracer [^11^C]PIB (Lockhart et al., 2007; Hellström-Lindahl et al., 2014). Such a large animal model is valuable in addition to rodent studies because of the pig’s larger and gyrated brain. We show that when the pig brain is injected with synthetic proteins or brain homogenates, the blood-brain barrier remains intact, the injected region’s protein levels are comparable to the characteristics in the human brain, and the in vivo binding characteristics allow for realistic quantification.

We validated our acute model by injecting α-synuclein preformed fibrils, AD human brain homogenate, or DLB human brain homogenate in pigs’ mPFC and visualized these regions *in vivo* using [^11^C]PIB PET. [^11^C]PIB uptake in the injection site was used as a proof of concept for this model. We found high regional [^11^C]PIB uptake in the AD homogenate and moderate uptake in α-synuclein preformed fibril injected regions. We also confirmed absence of specific uptake or binding of the radioligand in DLB homogenate injected or saline injected regions. Collectively, these results suggest that the model provides a tool for preclinical characterization of novel radioligands, including collecting information about the pharmacokinetics and affinities of the brain pathology.

[^11^C]PIB is a well-characterized radioligand for amyloid-β plaques (Price et al., 2005; Peretti et al., 2019), routinely used to quantify amyloid-β plaques and for differential diagnosis and staging in neurodegenerative diseases. Although [^11^C]PIB has the highest affinity to amyloid plaques, it also displays affinity towards other β-sheet structures like tau and α-synuclein. We chose to use [^11^C]PIB as proof of concept since it shows affinity to α-synuclein preformed fibrils (Fodero-Tavoletti et al., 2007; Ye et al., 2008) and AD brain homogenates (Klunk et al., 2004; Lockhart et al., 2007). By contrast, and as a negative control, [^11^C]PIB has no affinity to Lewy bodies commonly seen in PD or DLB histology (Fodero-Tavoletti et al., 2007; Ye et al., 2008), which we also confirmed both in vivo and in vitro.

To the best of our knowledge, this is the first time [^11^C]PIB has been tested in pigs with full arterial blood sampling and kinetic modeling. Our laboratory and Aarhus University (Alstrup et al., 2018) have previously performed [^11^C]PIB scans in pigs (unpublished), where the data was quantified using reference tissue modeling. Invasive kinetic modeling with [^11^C]PIB was challenging since the 1-tissue compartment model yielded a poor fit, while the 2-tissue compartment model failed, most likely because of the very fast metabolism of the parent compound. Instead, we used the graphical method, i.e., Logan Graphical analysis. We also used the SRTM2 with the occipital cortex as a reference region (Yaqub et al., 2008; Tolboom et al., 2009). In humans, SRTM2 modeling of [^11^C]PIB is commonly used with the cerebellum as the reference region, but when that was attempted in the pig brain, we got negative BP_ND_ values in DLB injected, saline injected and occipital cortex. Hence, we used the occipital cortex instead as a reference region.

Postmortem human brain homogenates from patients with relevant neurodegenerative disorders were introduced to “humanize” the pig model. We evaluated that the *B*_max_ in the injected pig brain was realistically representing what is seen in the individuals with AD who served as the donors of tissue homogenate. The observation that we found slightly lower *B*_max_ values in pig brain slices representing the AD homogenate injected regions compared to human brain slices from AD patients confirms the suitability of our pig model. We also performed a [^3^H]PIB saturation assay on the α-synuclein preformed fibril injected pig brain slices. Compared to the AD homogenate slices, the α-synuclein preformed fibril injected pig brain slices had a 2.4-times lower BP_ND_, as we also found in the in vivo PET studies. It can be argued that injection of human brain homogenates provides a more realistic model of the human AD brain compared to synthetic protein injections with, e.g., preformed fibrils but in any instance, the synthesized protein must be thoroughly evaluated in vitro before using it the model. In the current study, we used the highest available concentration of all the injectates for proof of concept. As an added value of the model in future studies, the concentration of the injectates can be varied to confirm the expected dose-dependent effect of radioligand binding.

Whereas strategy of intracerebrally injecting α-synuclein (and amyloid-β) and scanning animals immediately after previously has been used in rodents (Verdurand et al., 2018; Kuebler et al., 2020), this is the first study to involve larger animals. A few other large animal models of α-synuclein pathology have been published: the viral-vector model in minipigs (Lillethorup et al., 2018) and nonhuman primates (Kirik et al., 2002; Yang et al., 2015; Koprich et al., 2016), and α-synuclein protein or homogenate inoculation models also in nonhuman primates (Recasens et al., 2014; Shimozawa et al., 2017). The disadvantages of these models are that they are challenging to create, expensive to maintain and it often take months to develop pathology. National regulations on ethical considerations can also restrict access to experimental studies in nonhuman primates. By contrast, our model combines surgery and scanning on the same day, using non-survival pigs and a systematic scanning technique for in vivo radioligand characterization (Ettrup et al., 2013; Andersen et al., 2015; Jørgensen et al., 2018). Studies in pigs are cheaper than other large animals as the use of pigs is considered less ethically challenging.

Conventional strategies for intracerebral injection involve an MR-guided stereotactic approach (Glud et al., 2011). This requires MR-guided calculation of the stereotactic coordinates prior to surgery for the injection, which is a tedious and time-consuming procedure. In the present study, we employed a minimally invasive approach with a modified stereotactic instrument and a previously validated target point in the grey matter of mPFC (Jørgensen et al., 2017, 2018), which made the procedure much faster; the process including injection of the experimental substrates in mPFC was completed within 3-4 hours. The concern whether the blood-brain barrier would retain its integrity right after the intracerebral injection was addressed by the finding that the gadolinium-enhanced post-injection MR assured no blood-brain barrier leakage, except in the cases where the needle had induced minor traumatic hemorrhage – this was clearly outside the region with pathology. This observation was further supported by the saline-injected region having a radioligand uptake similar to the reference regions (Table 2).

Some limitations with the model presented should be mentioned. Although this model can be used for survival studies, we have only validated bilateral injection sites in the medial prefrontal cortex. A thorough in vitro evaluation of the proteins is necessary before commencing in vivo experiments, preferably including autoradiography with the radioligand to be evaluated. The latter includes identification of *K*_D_ and *B*_max_ to establish the in vitro binding potential which should reflect the PET signal. Further, the injection site constitutes a relatively small volume of interest which inherently is prone to noisy time-activity curves or to partial volume effect. Further, bleeding from dura or the cerebral tissue resulting from the injection could potentially impact the PET signal. We saw confined hematomas in 1 out of 5 injections but this was clearly recognized and when taken into account, it did not prevent a proper analysis. For future use of the model, we recommend to use hybrid PET/CT or PET/MR so that eventual hemorrhage can be accounted for.

## 5 Conclusions

We here provide a novel large model for assessment of novel radioligands targeting the brain and show its suitability for testing radioligands for brain regional proteinopathies. The large pig brain makes it suitable for neurosurgical procedures and the pigs can undergo multiple PET scans and frequent blood sampling. The described pig model represents a robust and efficient set-up with few limitations. The availability of a large animal α-synuclein model is instrumental for testing novel radioligands, not only for α-synuclein neuroimaging but also for other target proteins where the target is not naturally occurring in the brain, or where the presence can be artificially enhanced locally in the brain.

## 6 Acknowledgments

This project has received funding from the European Union’s Horizon 2020 research and innovation program under the Marie Skłodowska-Curie grant agreement No. 813528. This project also received funding from Parkinson foreningen, Denmark (R16-A247). Pontus Plavén Sigray was supported by the Lundbeck Foundation (R303-2018-3263). Natalie Beschorner was supported by the Lundbeck Foundation (R322-2019-2744). We would like to express our sincere gratitude to Patrick MacDonald Fisher for his technical assistance in MRI scanning protocols and data processing. We want to thank Lundbeck A/S, Valby, Denmark, for providing the α-synuclein preformed fibrils. This research project received human brain tissue from the Neuropathology Core of the Emory Center for Neurodegenerative Disease; we are grateful for their support. We would sincerely like to thank the staff and veterinarians at EMED, Panum, København University.

## 7 Author Contributions

NRR, LMJ, GMK: conceptualization and design. NRR, LMJ: surgical setup. NRR, AN, CAM, NB, EEB, SL: experimental studies. NRR, AN, CS, SL, PPS, MJ: analysis and software. NRR, GMK: resources. NR, PPS, LMJ, GMK: data curation. NRR: preparation of manuscript draft including figures. NRR, AN, CAM, NB, EEB, MJ, SL, MPP, CS, PPS, LMJ, GMK: manuscript review and editing. MP, CS, PPS, LMJ, GMK: supervision. NRR, MP, GMK: funding acquisition. All authors have read and agreed to the published version of the manuscript.

## 9 Data availability statement

All data, including MATLAB and R scripts, is available at a GitHub repository (https://github.com/nakulrrraval/Protien_inj_pig_model_PIB). All other requests are directed to this article’s corresponding or first author.

## References

Alstrup, A. K. O., Munk, O. L., and Landau, A. M. (2018). PET radioligand injection for pig neuroimaging. of Laboratory Animal … 44. Available at: http://sjlas.org/index.php/SJLAS/article/view/509.

Braak, H., and Braak, E. (2000). Pathoanatomy of Parkinson’s disease. J. Neurol. 247 Suppl 2, II3–10. doi:10.1007/PL00007758.

Capotosti, F., Vokali, E., Molette, J., Tsika, E., Ravache, M., Juergens, T., et al. (2020). Developing a novel alpha-synuclein positron emission tomography (PET) tracer for the diagnosis of synucleinopathies. Alzheimers. Dement. 16. doi:10.1002/alz.043249.

Donovan, L. L., Magnussen, J. H., Dyssegaard, A., Lehel, S., Hooker, J. M., Knudsen, G. M., et al. (2020). Imaging HDACs In Vivo: Cross-Validation of the [ 11 C]Martinostat Radioligand in the Pig Brain. Mol. Imaging Biol. 22, 569–577. doi:10.1007/s11307-019-01403-9.

Dunn, O. J. (1964). Multiple Comparisons Using Rank Sums. Technometrics 6, 241–252. doi:10.1080/00401706.1964.10490181.

Ettrup, A., Holm, S., Hansen, M., Wasim, M., Santini, M. A., Palner, M., et al. (2013). Preclinical safety assessment of the 5-HT2A receptor agonist PET radioligand [11C]cimbi-36. Mol. Imaging Biol. 15, 376–383. doi:10.1007/s11307-012-0609-4.

Fodero-Tavoletti, M. T., Smith, D. P., McLean, C. A., Adlard, P. A., Barnham, K. J., Foster, L. E., et al. (2007). In vitro characterization of Pittsburgh compound-B binding to Lewy bodies. J. Neurosci. 27, 10365–10371. doi:10.1523/JNEUROSCI.0630-07.2007.

Gillings, N. (2009). A restricted access material for rapid analysis of [(11)C]-labeled radiopharmaceuticals and their metabolites in plasma. Nucl. Med. Biol. 36, 961–965. doi:10.1016/j.nucmedbio.2009.07.004.

Glud, A. N., Hedegaard, C., Nielsen, M. S., Søorensen, J. C., Bendixen, C., Jensen, P. H., et al. (2011). Direct MRI-guided stereotaxic viral mediated gene transfer of alpha-synuclein in the Göttingen minipig CNS. Acta Neurobiol. Exp. 71, 508–518. Available at: https://www.ncbi.nlm.nih.gov/pubmed/22237496.

Hansen, H. D., Herth, M. M., Ettrup, A., Andersen, V. L., Lehel, S., Dyssegaard, A., et al. (2014). Radiosynthesis and in vivo evaluation of novel radioligands for pet imaging of cerebral 5-ht7 receptors. J. Nucl. Med. 55, 640–646. doi:10.2967/jnumed.113.128983.

Harding, J. D. (2017). Nonhuman Primates and Translational Research: Progress, Opportunities, and Challenges. ILAR J. 58, 141–150. doi:10.1093/ilar/ilx033.

Hellström-Lindahl, E., Westermark, P., Antoni, G., and Estrada, S. (2014). In vitro binding of [^3^H]PIB to human amyloid deposits of different types. Amyloid 21, 21–27. doi:10.3109/13506129.2013.860895.

Hooshyar Yousefi, B., Shi, K., Arzberger, T., Wester, H. J., Schwaiger, M., Yakushev, I., et al. (2019). Translational study of a novel alpha-synuclein PET tracer designed for first-in-human investigating. in NuklearMedizin 2019 (Georg Thieme Verlag KG), L25. doi:10.1055/s-0039-1683494.

Jørgensen, L. M., Baandrup, A. O., Mandeville, J., Glud, A. N., Sørensen, J. C. H., Weikop, P., et al. (2021). An FMRI-compatible system for targeted electrical stimulation. Research Square. doi:10.21203/rs.3.rs-313183/v1.

Jørgensen, L. M., Weikop, P., Svarer, C., Feng, L., Keller, S. H., and Knudsen, G. M. (2018). Cerebral serotonin release correlates with [11C]AZ10419369 PET measures of 5-HT1B receptor binding in the pig brain. J. Cereb. Blood Flow Metab. 38, 1243–1252. doi:10.1177/0271678X17719390.

Jørgensen, L. M., Weikop, P., Villadsen, J., Visnapuu, T., Ettrup, A., Hansen, H. D., et al. (2017). Cerebral 5-HT release correlates with [11C]Cimbi36 PET measures of 5-HT2A receptor occupancy in the pig brain. J. Cereb. Blood Flow Metab. 37, 425–434. doi:10.1177/0271678X16629483.

Kaide, S., Watanabe, H., Shimizu, Y., Iikuni, S., Nakamoto, Y., Hasegawa, M., et al. (2020). Identification and Evaluation of Bisquinoline Scaffold as a New Candidate for α-Synuclein-PET imaging. ACS Chem. Neurosci. 11, 4254–4261. doi:10.1021/acschemneuro.0c00523.

Keller, S. H., Svarer, C., and Sibomana, M. (2013). Attenuation correction for the HRRT PET-scanner using transmission scatter correction and total variation regularization. IEEE Trans. Med. Imaging 32, 1611–1621. doi:10.1109/TMI.2013.2261313.

Kirik, D., Rosenblad, C., Burger, C., Lundberg, C., Johansen, T. E., Muzyczka, N., et al. (2002). Parkinson-Like Neurodegeneration Induced by Targeted Overexpression of α-Synuclein in the Nigrostriatal System. Journal of Neuroscience 22, 2780–2791. doi:10.1523/JNEUROSCI.22-07-02780.2002.

Klunk, W. E., Engler, H., Nordberg, A., Wang, Y., Blomqvist, G., Holt, D. P., et al. (2004). Imaging brain amyloid in Alzheimer’s disease with Pittsburgh Compound-B. Ann. Neurol. 55, 306–319. doi:10.1002/ana.20009.

Koprich, J. B., Johnston, T. H., Reyes, G., and Omana, V. (2016). Towards a nonhuman primate model of alpha-synucleinopathy for development of therapeutics for Parkinson’s disease: optimization of AAV1/2 delivery …. PLoS. Available at: https://journals.plos.org/plosone/article?id=10.1371/journal.pone.0167235.

Korat, Š., Bidesi, N. S. R., Bonanno, F., Di Nanni, A., Hoàng, A. N. N., Herfert, K., et al. (2021). Alpha-Synuclein PET Tracer Development—An Overview about Current Efforts. Pharmaceuticals 14, 847. doi:10.3390/ph14090847.

Kuebler, L., Buss, S., Leonov, A., Ryazanov, S., Schmidt, F., Maurer, A., et al. (2020). [11C]MODAG-001—towards a PET tracer targeting α-synuclein aggregates. Eur. J. Nucl. Med. Mol. Imaging. doi:10.1007/s00259-020-05133-x.

Lashuel, H. A., Overk, C. R., Oueslati, A., and Masliah, E. (2013). The many faces of α-synuclein: from structure and toxicity to therapeutic target. Nat. Rev. Neurosci. 14, 38–48. doi:10.1038/nrn3406.

Lázaro, D. F., Bellucci, A., Brundin, P., and Outeiro, T. F. (2019). Editorial: Protein Misfolding and Spreading Pathology in Neurodegenerative Diseases. Front. Mol. Neurosci. 12, 312. doi:10.3389/fnmol.2019.00312.

Lillethorup, T. P., Glud, A. N., Landeck, N., Alstrup, A. K. O., Jakobsen, S., Vang, K., et al. (2018). In vivo quantification of glial activation in minipigs overexpressing human α-synuclein. Synapse 72, e22060. doi:10.1002/syn.22060.

Li, X., Morgan, P. S., Ashburner, J., Smith, J., and Rorden, C. (2016). The first step for neuroimaging data analysis: DICOM to NIfTI conversion. J. Neurosci. Methods 264, 47–56. doi:10.1016/j.jneumeth.2016.03.001.

Lockhart, A., Lamb, J. R., Osredkar, T., Sue, L. I., Joyce, J. N., Ye, L., et al. (2007). PIB is a non-specific imaging marker of amyloid-beta (Abeta) peptide-related cerebral amyloidosis. Brain 130, 2607–2615. doi:10.1093/brain/awm191.

Logan, J., Fowler, J. S., Volkow, N. D., Wolf, A. P., Dewey, S. L., Schlyer, D. J., et al. (1990). Graphical analysis of reversible radioligand binding from time-activity measurements applied to [N-11C-methyl]-(-)-cocaine PET studies in human subjects. J. Cereb. Blood Flow Metab. 10, 740–747. doi:10.1038/jcbfm.1990.127.

Makky, A., Bousset, L., Polesel-Maris, J., and Melki, R. (2016). Nanomechanical properties of distinct fibrillar polymorphs of the protein α-synuclein. Sci. Rep. 6, 37970. doi:10.1038/srep37970.

Matheson, G. J. (2019). kinfitr: Reproducible PET Pharmacokinetic Modelling in R. bioRxiv, 755751. doi:10.1101/755751.

Mathis, C. A., Lopresti, B. J., Ikonomovic, M. D., and Klunk, W. E. (2017). Small-molecule PET Tracers for Imaging Proteinopathies. Semin. Nucl. Med. 47, 553–575. doi:10.1053/j.semnuclmed.2017.06.003.

Parker, C. A., Gunn, R. N., Rabiner, E. A., Slifstein, M., Comley, R., Salinas, C., et al. (2012). Radiosynthesis and characterization of 11C-GSK215083 as a PET radioligand for the 5-HT6 receptor. J. Nucl. Med. 53, 295–303. doi:10.2967/jnumed.111.093419.

Peretti, D. E., Reesink, F. E., Doorduin, J., de Jong, B. M., De Deyn, P. P., Dierckx, R. A. J. O., et al. (2019). Optimization of the k2′ Parameter Estimation for the Pharmacokinetic Modeling of Dynamic PIB PET Scans Using SRTM2. Frontiers in Physics 7, 212. doi:10.3389/fphy.2019.00212.

Price, J. C., Klunk, W. E., Lopresti, B. J., Lu, X., Hoge, J. A., Ziolko, S. K., et al. (2005). Kinetic modeling of amyloid binding in humans using PET imaging and Pittsburgh Compound-B. J. Cereb. Blood Flow Metab. 25, 1528–1547. doi:10.1038/sj.jcbfm.9600146.

Recasens, A., Dehay, B., Bové, J., Carballo-Carbajal, I., Dovero, S., Pérez-Villalba, A., et al. (2014). Lewy body extracts from Parkinson disease brains trigger α-synuclein pathology and neurodegeneration in mice and monkeys. Ann. Neurol. 75, 351–362. doi:10.1002/ana.24066.

Saikali, S., Meurice, P., Sauleau, P., Eliat, P.-A., Bellaud, P., Randuineau, G., et al. (2010). A three-dimensional digital segmented and deformable brain atlas of the domestic pig. J. Neurosci. Methods 192, 102–109. doi:10.1016/j.jneumeth.2010.07.041.

Schneider, C. A., Rasband, W. S., and Eliceiri, K. W. (2012). NIH Image to ImageJ: 25 years of Image Analysis HHS Public Access.

Shalgunov, V., Xiong, M., L’Estrade, E. T., Raval, N. R., Andersen, I. V., Edgar, F. G., et al. (2020). Blocking of efflux transporters in rats improves translational validation of brain radioligands. EJNMMI Res. 10, 124. doi:10.1186/s13550-020-00718-x.

Shimozawa, A., Ono, M., Takahara, D., Tarutani, A., Imura, S., Masuda-Suzukake, M., et al. (2017). Propagation of pathological α-synuclein in marmoset brain. Acta Neuropathol Commun 5, 12. doi:10.1186/s40478-017-0413-0.

Sureau, F. C., Reader, A. J., Comtat, C., Leroy, C., Ribeiro, M. J., Buvat, I., et al. (2008). Impact of image-space resolution modeling for studies with the high-resolution research tomograph. J. Nucl. Med. 49, 1000–1008. doi:10.2967/jnumed.107.045351.

Tjerkaski, J., Cervenka, S., Farde, L., and Matheson, G. J. (2020). Kinfitr - an open-source tool for reproducible PET modelling: validation and evaluation of test-retest reliability. EJNMMI Res. 10, 77. doi:10.1186/s13550-020-00664-8.

Tolboom, N., Yaqub, M., Boellaard, R., Luurtsema, G., Windhorst, A. D., Scheltens, P., et al. (2009). Test-retest variability of quantitative [11C]PIB studies in Alzheimer’s disease. Eur. J. Nucl. Med. Mol. Imaging 36, 1629–1638. doi:10.1007/s00259-009-1129-6.

Verdurand, M., Levigoureux, E., Zeinyeh, W., Berthier, L., Mendjel-Herda, M., Cadarossanesaib, F., et al. (2018). In Silico, in Vitro, and in Vivo Evaluation of New Candidates for α-Synuclein PET Imaging. Mol. Pharm. 15, 3153–3166. doi:10.1021/acs.molpharmaceut.8b00229.

Villadsen, J., Hansen, H. D., Jørgensen, L. M., Keller, S. H., Andersen, F. L., Petersen, I. N., et al. (2017). Automatic delineation of brain regions on MRI and PET images from the pig. J. Neurosci. Methods 294, 51–58. doi:10.1016/j.jneumeth.2017.11.008.

Winterdahl, M., Audrain, H., Landau, A. M., Smith, D. F., Bonaventure, P., Shoblock, J. R., et al. (2014). PET brain imaging of neuropeptide Y2 receptors using N-11C-methyl-JNJ-31020028 in pigs. J. Nucl. Med. 55, 635–639. doi:10.2967/jnumed.113.125351.

Yang, W., Wang, G., Wang, C.-E., Guo, X., Yin, P., Gao, J., et al. (2015). Mutant Alpha-Synuclein Causes Age-Dependent Neuropathology in Monkey Brain. Journal of Neuroscience 35, 8345–8358. doi:10.1523/JNEUROSCI.0772-15.2015.

Yaqub, M., Tolboom, N., Boellaard, R., van Berckel, B. N. M., van Tilburg, E. W., Luurtsema, G., et al. (2008). Simplified parametric methods for [11C]PIB studies. Neuroimage 42, 76–86. doi:10.1016/j.neuroimage.2008.04.251.

Ye, L., Velasco, A., Fraser, G., Beach, T. G., Sue, L., Osredkar, T., et al. (2008). In vitro high affinity alpha-synuclein binding sites for the amyloid imaging agent PIB are not matched by binding to Lewy bodies in postmortem human brain. J. Neurochem. 105, 1428–1437. doi:10.1111/j.1471-4159.2008.05245.x.

Lammertsma, A. A., and Hume, S. P. (1996). Simplified reference tissue model for PET receptor studies. Neuroimage 4, 153–158. doi:10.1006/nimg.1996.0066.

