## Supplementary materials for "An *in vivo* pig model for testing novel PET radioligands targeting cerebral protein aggregates"

Supplementary Material

### 1 α-synuclein preformed fibrils and post-mortem human brain pathological details

α-synuclein preformed fibrils were produced at Lundbeck A/S, Denmark, using their optimized protocols following previously published literature (Makky et al., 2016). Preformed fibrils were sonicated after fibrillization, aliquoted, and stored at −80°C until further use.

The post-mortem human brain homogenates used are a mixture of two regions (frontal and temporal) for each of two different subjects with each disease (i.e., AD and DLB). Samples, predominantly from the gray matter, were weighed from all regions and subjects; approximately 100 mg was used. All samples (both diseases separately) were added to an Eppendorf tube followed by an adequate amount of saline to make a 10% solution (w/v) and homogenized using an automatic tissue homogenizer (IKA® T10basic ULTRA-TURRAX® Disperser, IKA®-Werke GmbH & Co. KG, Staufen, Germany) for approximately 3 min to make a homogenous solution, aliquoted and stored at −80°C until further use.

**Supplementary Table 1.** Neuropathological characteristics of the post-mortem human brains used in the study. Details include primary neuropathological diagnosis: Alzheimer's disease (AD) or Dementia with Lewy bodies-Neocortical (LBD), frequency of Lewy bodies: frequent (F) or sparse to moderate (S-P), Braak staging, ABC, CERAD (The Consortium to Establish a Registry for Alzheimer's Disease), Thal staging, Post-mortem interval (PMI), age at onset, age at death, ApoE (Apolipoprotein E) genotype and sex of the brain available for this study.

| **Primary Diagnosis** | **Frontal Lewy Bodies** | **Parietal Lewy Bodies** | **Braak Stage** | **ABC** | **CERAD** | **Thal** | **PMI (h)** | **Age at onset (y)** | **Age at Death (y)** | **ApoE gene** | **Sex** |
| --- | --- | --- | --- | --- | --- | --- | --- | --- | --- | --- | --- |
| **AD1** | 0 | 0 | VI | 3 | 3 | 5 | 11 | 53 | 70 | E3/E4 | m |
| **AD2** | 0 | 0 | VI | 3 | 3 | 5 | 14.5 | 60 | 68 | E3/E3 | m |
| **LBD1** | F | S-M | II | 0 | 0 | 0 | 15.5 | 47 | 68 | E3/E3 | m |
| **LBD2** | F | S-M | II | 0 | 0 | 0 | 19.5 | 61 | 69 | E3/E3 | m |

### 2 Images and diagrams of the surgical procedure

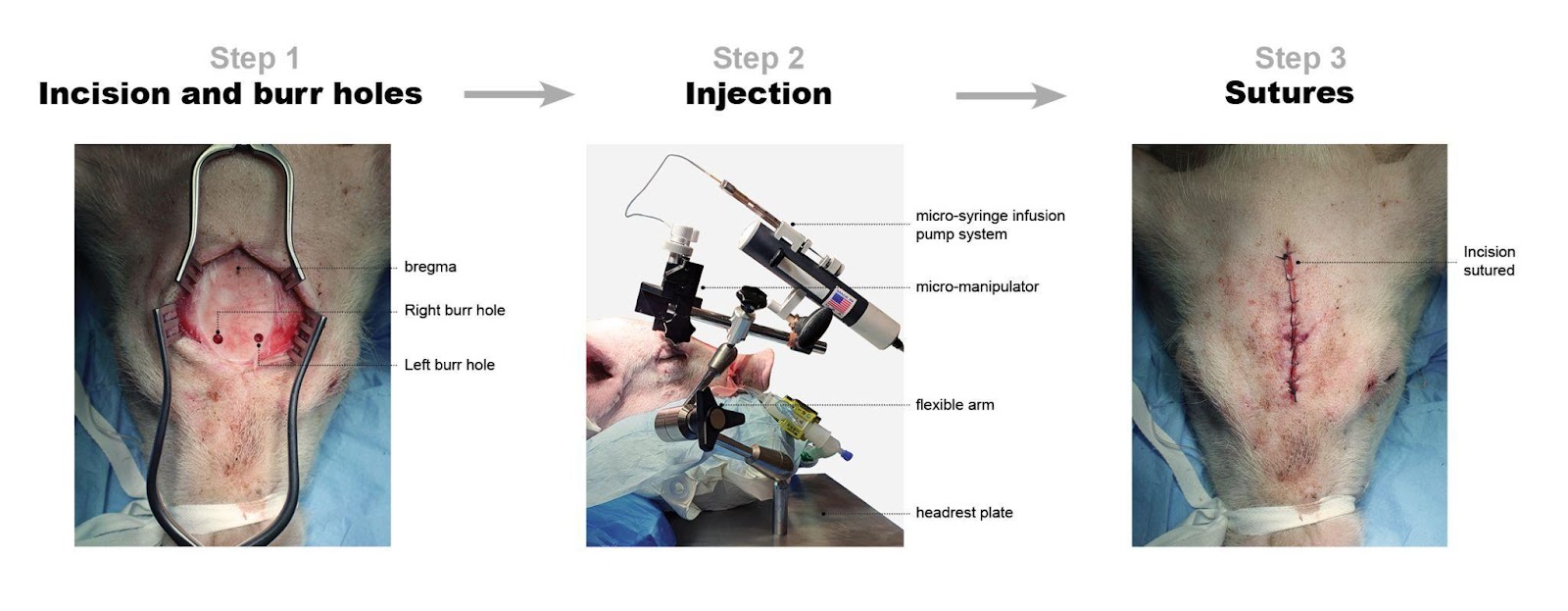

**Supplementary Figure 1.** Surgical procedure. *Step 1:* Incision and burr holes. Sagittal midline incision followed by the location of bregma. Burr holes are placed in precise relation to the bregma as per protocol. *Step 2:* Injection of substrates. The image shows an injection procedure and the details of the modified stereotactic frame. *Step 3:* Sutures. Sutures are placed, and the animal is ready for transport to the scanners.

### 3 [^11^C]PIB radiochemistry

[^11^C]PIB was prepared by reacting [^11^C]methyl triflate with 1.0 mg nor-PIB precursor dissolved in 300 mL methylethylketone (MEK) for 3 min at 80°C (Supplementary Figure 2). The reaction mixture was diluted with 4.5 mL 0.1% phosphoric acid and subsequently injected onto an Onyx Monolithic C18 semi-preparative HPLC column (10 x 100 mm, Phenomenex, Torrance, CA, USA). Eluent: 10 mM ascorbic acid in 0.1% phosphoric acid / ethanol 96% (70:30); flow rate 6.0 mL/min; wavelength l=250 nm. The fraction containing the product was collected within 1 min (6.0 mL) and passed through a sterile filter into a 20 mL sterile vial containing 9.0 mL 0.1 M phosphate buffer (pH=7), giving a final solution for injection containing ≤10%(v/v) ethanol.

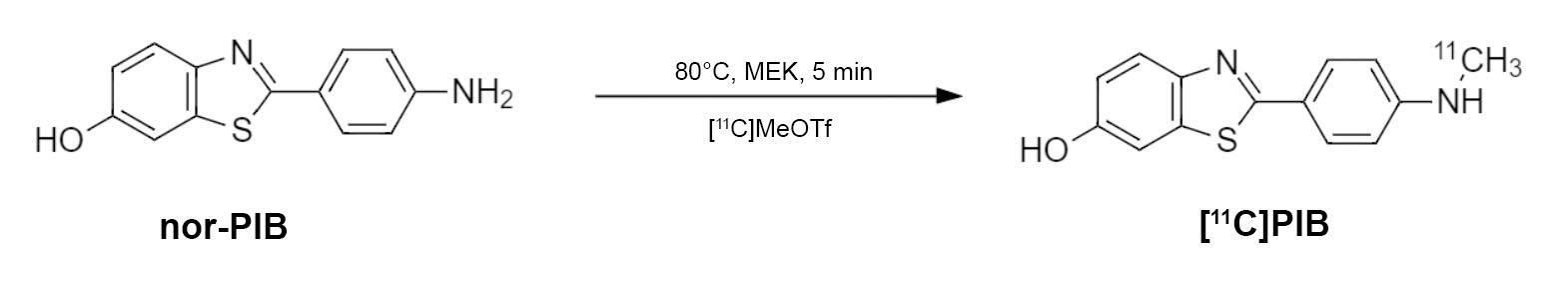

**Supplementary Figure 2.** Synthesis of [^11^C]PIB.

### 4 Addition analysis with blood and plasma

To exclude the possibility that the fast metabolism of [^11^C]PIB was due to the instability of the compound in plasma or blood, we verified the stability of [^11^C]PIB. The samples were spiked with [^11^C]PIB and incubated at room temperature for up to 75 min. Samples were processed and analyzed by radio-HPLC as described above in the main manuscript. We found that at room temperature, [^11^C]PIB was completely stable in plasma and whole blood for up to 75 min of incubation.

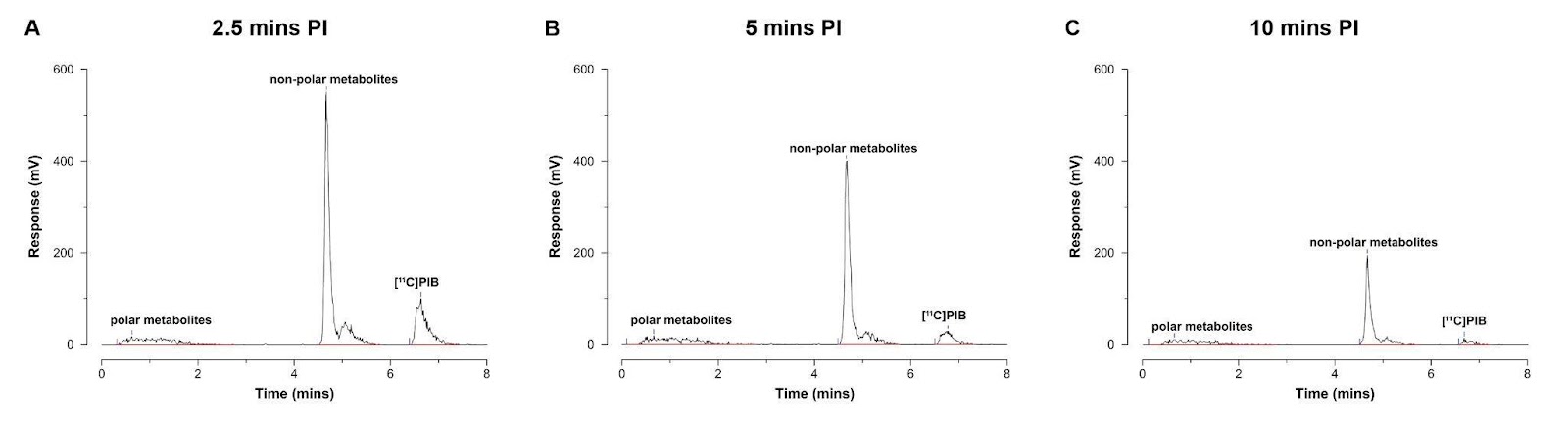

**Supplementary Figure 3.** HPLC chromatograms. A-C) Representative radio-chromatograms 2.5 min (A), 5 min (B), and 10 min post-injection of [^11^C]PIB (n=1).

### 5 Fluorescence immunostaining

Fluorescence immunostaining was performed on sections containing the injection site to validate the injection site and confirm injection of injectate. The sections were processed for standard immunohistochemistry (IHC) with α-synuclein (GTX21904-Mouse [4B12], GeneTex, Hsinchu City, Taiwan) and amyloid-ꞵ (ab252816-Rat [2E9], Abcam, Cambridge, UK) primary antibodies detected with goat anti-mouse or goat anti-rat IgG H&L (Alexa Fluor® 647) (ab150115/ab150167, Abcam, Cambridge, UK) secondary antibodies. The frozen sections were first fixed in 4% formaldehyde for 20 min. After a 10 min wash in phosphate-buffered saline (PBS), antigen retrieval was performed in a microwave oven in 25 mM sodium citrate buffer pH 7.5 for α-synuclein containing section or Tris/EDTA buffer pH 9.0 for amyloid-ꞵ containing section. Buffer temperature was raised to boiling for 5 secs, sections left in the microwave for 10 min, then 20 min under the hood. After, they were washed with PBS-TritonX100 0.4% and then incubated for 60 min with PBS + 0.4% TritonX100 and 5% bovine serum albumin (BSA). Buffer was poured off, sectioned for incubated overnight in primary antibody (α-synuclein: 0.2 ng/ml) (amyloid-ꞵ: 0.5 µg/ml) in PBS + 0.1% Tween20 at 4 °C. The next day, sectioned were washed thrice in PBS and then incubated in secondary antibody (diluted 1:200) in PBS + 0.1% Tween20 for 1 hour at room temperature. Finally, the sections were washed thrice in PBS followed by one wash in deionized H_2_O (dH_2_O). After the standard IHC, the sections were washed in increasing ethanol concentration (1 min in 70% ethanol and 1 min in 85%) and then incubated in 0.25% thioflavin S solution in 85% ethanol (filtered) for 15 mins. Incubation was followed by a wash in decreasing ethanol concentration and finally a wash with dH_2_O. After the staining protocol, the sections were overnight mounted in EverBrite™ Hardset Mounting Medium (Biotium, Inc., Fremont, CA, USA). Sections were imaged using an EC Plan-Neofluoar 5x/0.16 objective on an Axio Observer 7 fitted with a motorized stage and Axiocam 506mono CCD camera (Carl Zeiss, Birkerød, Denmark) to create stitched images covering large regions of interest. ThS was imaged using a filter set comprising a 470/40 nm bandpass for excitation, 495 nm beamsplitter, and 525/50 nm bandpass emission (Filter Set 38 HE, Carl Zeiss, Birkerød, Denmark). For Alexa Fluor® 647, an excitation of 640/30 nm, beamsplitter of 660, and emission of 690/50 filter set was used (Filter Set 50, Carl Zeiss, Birkerød, Denmark).

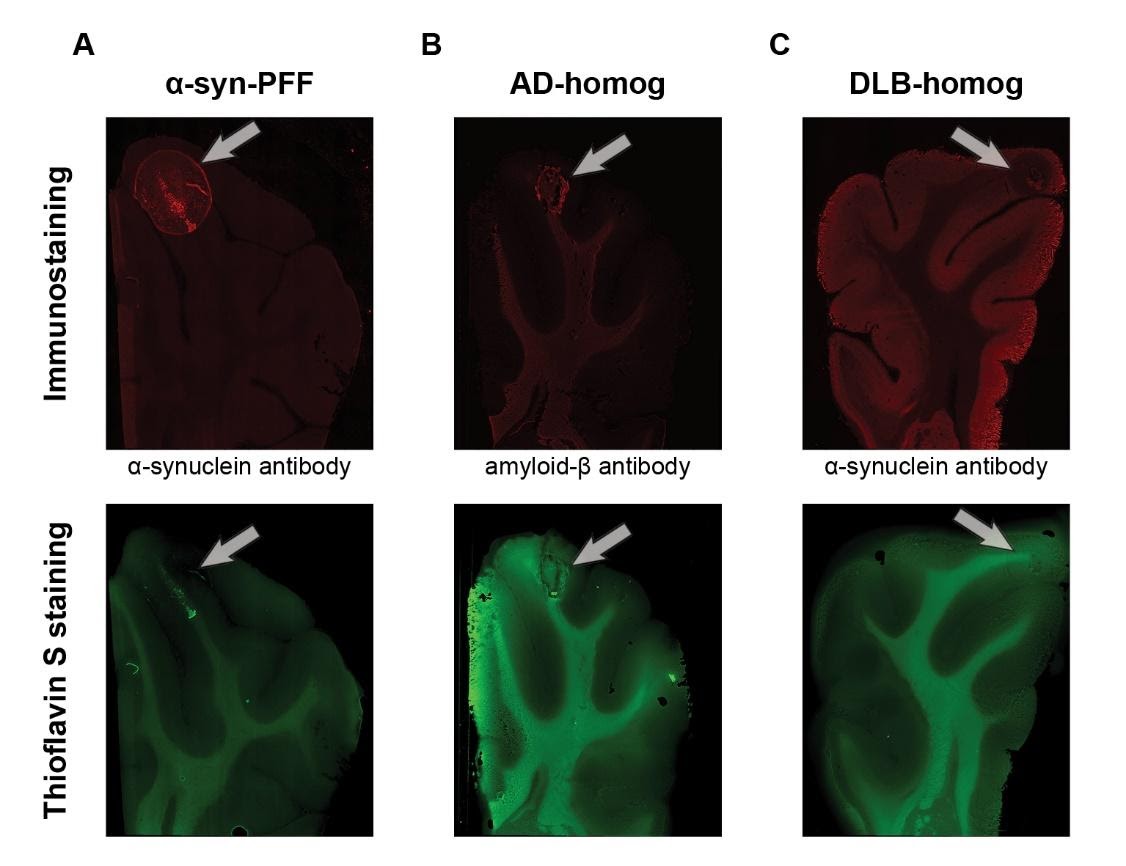

**Supplementary Figure 4.** Successful injections were ensured by immunostaining (top row) and thioflavin S staining (bottom row). Representative examples of A) α-synuclein-preformed-fibrils injected hemisphere, B) AD-homogenate injected hemisphere, and C) DLB-homogenate injected hemisphere.

### 6 [^3^H]PIB saturation assay

Sections were thawed to room temperature for 45 min before pre-washing twice for 10 min in assay buffer (PBS, 1% BSA, 10% ethanol, and pH 7.4). Sections were incubated in assay buffer containing varying concentrations of [^3^H]PIB (0 to 5 nM for AD-homogenate-injected pig brain and AD human brain slices) (0 to 40 nM for α-synuclein-preformed-fibril-injected pig brain slices) for total binding (TB) and the same varying concentrations of [^3^H]PIB with 100 µM of thioflavin S for non-specific binding (NSB). The sections were incubated for 60 min. Incubation was terminated by three 5-min washes with 4°C wash buffer (PBS, 10% ethanol, and pH 7.4) followed by a rapid rinse in 4°C dH_2_O.

After washing, the slides were rapidly air-dried and fixated overnight in a paraformaldehyde vapor chamber in cold storage (4°C). The next day, the samples were moved to an exicator for 60 min to remove any excess moisture and then placed in a cassette for autoradiography with tritium sensitive image plates (BAS-IP TR2040 E, Science Imaging Scandinavia AB, Nacka, Sweden) along with radioactive tritium standards (ART0123, American Radiolabeled Chemical, Inc., St. Loui, MO, USA). The image plates were exposed for three days. After the exposure, the image plates were read using an Amersham™ Typhoon™ IP (Cytiva, Uppsala, Sweden) at 10 µm resolution. Calibration, quantification, and data evaluation were done using ImageJ software (NIH Image, Bethesda, MD, USA) (Schneider et al., 2012). The regions of interest were hand-drawn. The four-parameter general curve fit (David Rodbard, NIH) of decay corrected tritium standards was used to convert mean pixel density (grayscale) to nCi/mg Tissue Equivalent (TE). TB was determined in the pathology-rich regions or at the injection site from TB slides, while NSB was determined in pathology-rich regions or at the injection site from NSB slides. Finally, the decay-corrected specific activity of [^3^H]PIB was used to convert nCi/mg TE to fmol/mg TE which was used to calculate the *B*_max_ and *K*_D_ and the binding potential (BP) as the ratio between the two parameters.

### 7 Kinetic Modelling and Population-based parent fraction

Supplementary Figure 5 shows the volume of interest for the injection regions that are added to the existing atlas and further used in the study. All animals' regional time-activity curves and blood data, including whole blood activity, plasma activity, and parent fraction throughout the scan, were loaded to *kinfitr*. The parent fraction curve was fit to a model. We tested multiple models for the parent fraction curve and found the inverted gamma function (inverse gamma model) to have the best fit using the Bayesian information criterion (data not shown). The blood volume (V_B_) was set to 0.05 (5%). Blood delay (arterial input function delay compared to brain time-activity curve) and weights were calculated and set according to standard protocol. The optimal threshold time (t*) was computed using the inbuilt function of *kinfitr* called "Logan_tstar." We found a t* of 10 (last 10 data points from the Logan plot) ideal since it had the least maximum percentage of variance and stable V_T_ values.

Due to a technical failure with the HPLC equipment, the parent fraction curve from the last three scans could not be estimated. Instead, we created a population-based parent fraction curve from the first four scans and used it for the last three scans (Table 1, Supplementary Figure 6). This was created by averaging the available parent fraction curves at their respective time-point. The population-based curve was validated by plotting Logan V_T_ values from the individual parent fraction curve against V_T_ values from the population-based parent fraction curve in the first four animals. We found a highly significant correlation between these two V_T_ values (Pearson r = 0.99, p < 0.0001) (Supplementary Figure 6).

For reference tissue modeling using the SRTM2, the k'_2_ was first estimated by fitting the time-activity curves from all animals to their occipital cortex using SRTM (Lammertsma and Hume, 1996). The k'_2_ parameter was extracted from all the high binding regions (AD-homogenate and ⍺-synuclein-preformed-fibril and injected regions) and averaged, providing a k'_2_ of 0.03. After fixing k'_2_ to 0.03, we fit the time-activity curves from all the animals to their occipital cortex using the SRTM2 to extract the non-displaceable binding potential (BP_ND_) values.

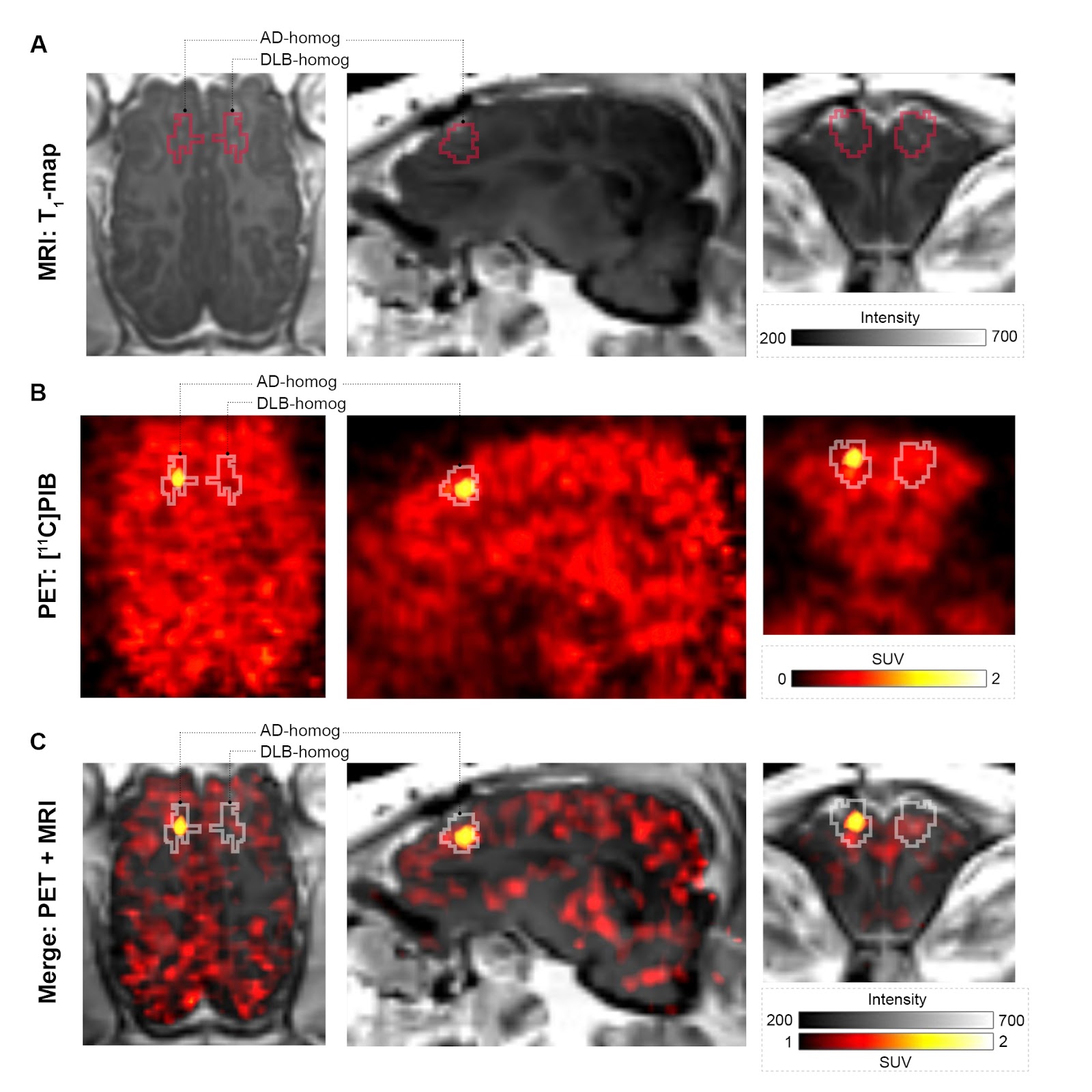

**Supplementary Figure 5.** Injection region volume of interests on representative A) T_1_-map MRI, B) [^11^C]PIB summed PET image, and C) merged PET + MR image in a pig injected with AD-homogenates and DLB-homogenates. PET image is created over the entire scan duration (0-90 mins) using the "Triangle" interpolation function of PMOD for better visualization of the injection region. The SUV scaling is changed for the merged image to reduce the background.

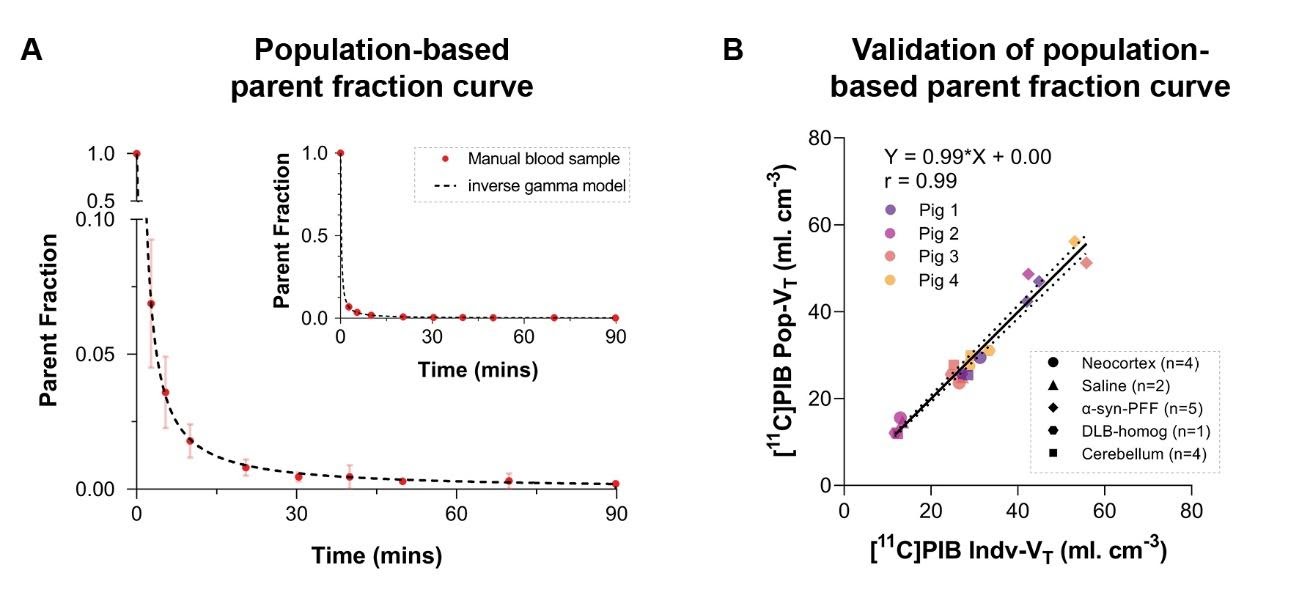

**Supplementary Figure 6**. Population-based parent fraction curve and its validation. A) Population-based parent fraction curves from n=4 animals fit an inverse gamma model. B) Validation of the population-based parent fraction curve. Logan V_T_ values calculated from the population-based parent fraction curve plotted against Logan plot V_T_ values calculated from individual parent fraction curves from the same animals. Equation from the linear regression and Pearson's r value is inserted.

### 8 Reference

Lammertsma, A. A., and Hume, S. P. (1996). Simplified reference tissue model for PET receptor studies. *Neuroimage* 4, 153–158. doi:10.1006/nimg.1996.0066.

Makky, A., Bousset, L., Polesel-Maris, J., and Melki, R. (2016). Nanomechanical properties of distinct fibrillar polymorphs of the protein α-synuclein. Sci. Rep. 6, 37970. doi:10.1038/srep37970.

Schneider, C. A., Rasband, W. S., and Eliceiri, K. W. (2012). NIH Image to ImageJ: 25 years of Image Analysis HHS Public Access.
